# A low-cost and open-source olfactometer to precisely deliver single odours and odour mixtures

**DOI:** 10.1101/2025.09.11.675563

**Authors:** Connor Doyle, Jieni Wang, Elisa Galliano, Chloé Guillaume

## Abstract

Despite its profound influence on human emotion and memory, and its critical importance in animal behaviour, olfaction has historically been understudied compared to other sensory modalities. Contributing to this relative neglect may be challenges inherent to odour delivery: unlike visual or auditory stimuli, odorants must be vaporised, transported, and actively cleared, requiring specialised equipment and creating temporal delays and contamination risks. While commercial olfactometers provide reliable solutions, they can be expensive and, due to proprietary software, may be difficult to customise. Existing custom designs, though excellent, often require specialized technical skills and/or access to engineering workshops. Thus, to help widen access to well-controlled odour delivery, we developed a low-cost, fully open-source olfactometer that can be assembled without specialised facilities or engineering skills and customised according to experimental needs. Our design prioritises affordability, accessibility, and versatility while maintaining temporal precision and stimulus control across applications from rodent behaviour to human psychophysics. This methods articles provides full design specifications, assembly details, and behavioural validation across both rodents and humans with the aim of reducing barriers and enabling more laboratories to study this crucial sensory system.

## Introduction

*“I remember my childhood names for grasses and secret flowers. I remember where a toad may live and what time the birds awaken in the summer - and what trees and seasons smelled like - how people looked and walked and smelled even. **The memory of odors is very rich**.” East of Eden, John Steinbeck*

Olfaction has long been regarded as a secondary sense, influenced perhaps by the persistent but mistaken belief that the human sense of smell is relatively weak compared with that of other animals (1). This misconception likely stems from the relatively small size of the human olfactory bulbs (in relation to the rest of the brain)(2), and the mistaken conflation of anatomical size and functional importance. In reality, the olfactory system, including that of humans, is a powerful modulator of memory, emotion and decision-making thanks to its direct projections to regions such as the limbic system and prefrontal cortex (3). Indeed, following the COVID-19 pandemic with high anosmia comorbidity, there has been an increase in both public and scientific awareness of the impact of olfaction for quality of life (4,5).

While the importance of olfaction in humans is often most apparent when it is lost, rodents, one of the most widely used model organisms in neuroscience, have always been known to rely heavily on olfaction for key behaviours including foraging, predator avoidance, mating, and parenting (6–8). Concurrently, the large and dorsally located rodent olfactory bulbs provide exceptional experimental accessibility for neuronal functional investigation. However, despite this combination of ethological importance and experimental tractability, olfactory research remains less developed than other sensory modalities (9,10). One factor contributing to this relative lack of study, might be a practical issue: the difficulty of precisely controlling olfactory stimuli. Unlike light or sound, which can be switched on and off almost instantaneously, odour molecules must be vaporized, transported through the air, and then actively cleared after delivery, to prevent odours lingering, creating delays and potential variability in stimulus timing (11,12). Additional challenges arise from the need to maintain stable airflow, detect odour concentration using specialised equipment such as photoionization detectors, and minimise contamination caused by adsorption of odorants to tubing and valves.

Despite these difficulties, numerous commercial and custom-built olfactometers exist that address many of these challenges. Commercial olfactometers provide reliable, ready-to-use solutions, but they are often expensive (in the tens of thousands of dollars range) and may be difficult to adapt to specific experimental needs due to the closed nature of their software. Many custom designs described in the literature are excellent, offering precise control, detailed construction guides, and thorough characterisation of their performance (13–20). However, none fully address all of our specific requirements: (a) affordable and scalable; (b) simple to assemble without specialised skills or access to mechanical/electronic workshops; (c) easily transportable across testing rooms or setups; (d) operated by fully open-source and customizable software; (e) suitable for a variety of experimental configurations, ranging from human psychophysics experiments, to rodent freely moving and head-fixed behavioural experiments; (f) suitable for multi-subject setups for simultaneous testing; (g) capable of delivering both single odours and mixtures while maintaining constant air pressure.

To address this, we have developed a low-cost, fully open-source olfactometer that can be assembled and operated without specialized workshops or extensive engineering expertise. Here we present detailed assembly instructions, which we hope will enable any researchers interested in building this device the necessary guidance to easily do so. Furthermore, we also provide olfactometric and behavioural validation of the olfactometer performance across a wide (although not exhaustive) list of essential variables, such as timing, tubing length, and air dilution. Finally, we present data from experiments performed using the olfactometer with both rodents and humans, as an example of the sort of experiments that may be performed using this olfactometer.

By providing these resources, we aim to lower the barriers to conducting olfactory research and enable more laboratories to explore the rich scientific questions that can be addressed in different models. Additionally, we hope that others can take our design as inspiration and adapt and extend it and share their improvements in keeping with the open-science ethos of this work.

## Results

### Overall design

Our olfactometer comprises three main components: (I) an air supply system, with corresponding pressure and flow regulation, (II) a modular odour system containing four odour modules, which may be expanded as desired, and (III) an electronic control system that regulates odour selection and timing via solenoid valves operated using open-source software on a standard desktop computer (Fig. 1A).

**Fig 1.**
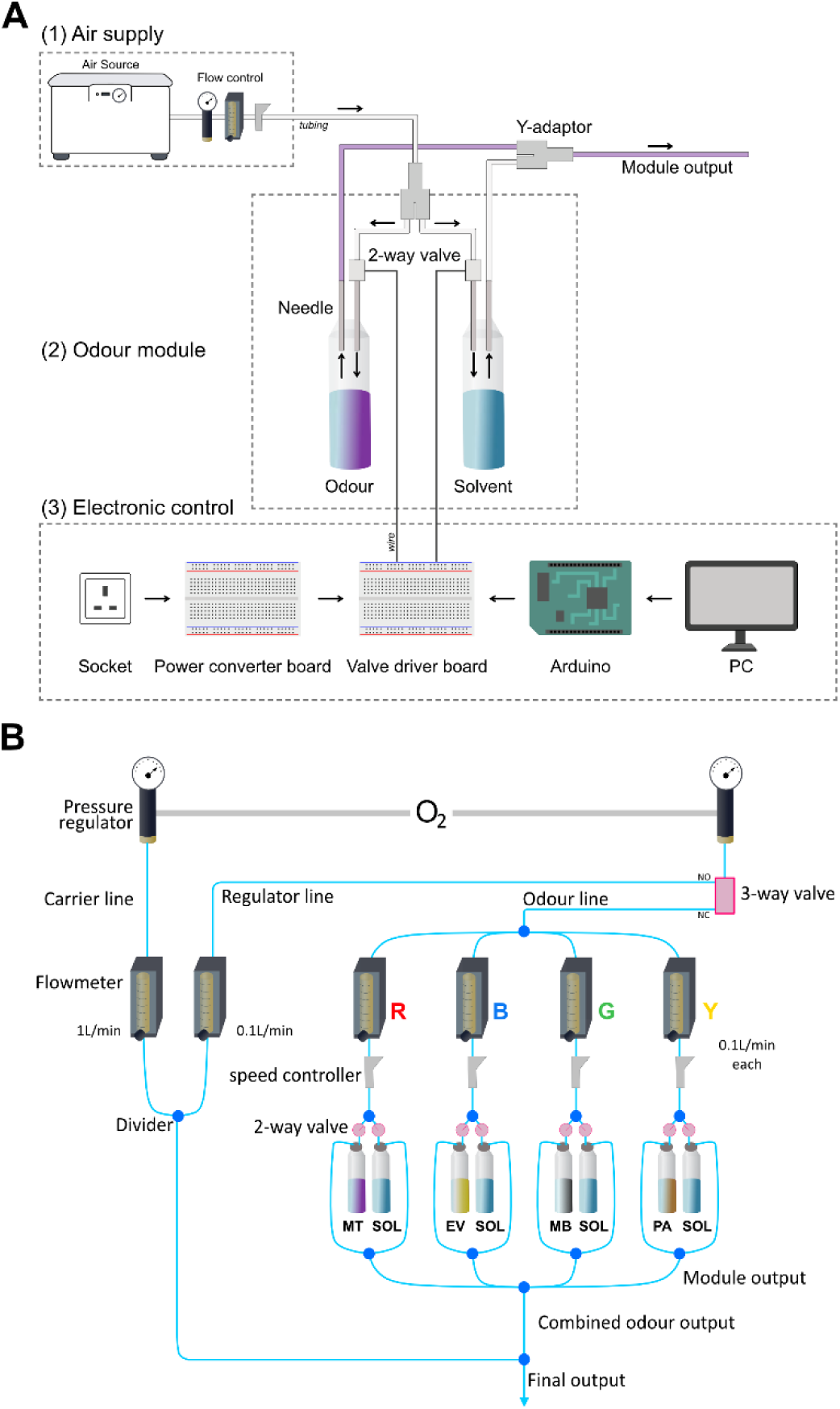
Olfactometer design and operation. **(A)** System overview showing the three main components: (1) air supply system, (2) detailed view of a single odour module showing the paired bottle design with air flow control systems, inlet/outlet needles, solenoid valve control, and (3) integration with the Arduino-based control system via valve driver boards and power converter boards. **(B)** Schematic representation of the air flow system across the four colour-coded odour modules (Red, Blue, Green, Yellow) each containing paired solvent (SOL) and odorant bottles, and electronic control via solenoid valves. The carrier stream (1 L/min) flows continuously to maintain stable baseline conditions, while odour streams (0.1 L/min each) pass through individual modules before merging at the output. Abbreviations: MT, methyl tiglate; EV, ethyl valerate; MB, methyl butyrate; PA, pentyl acetate; NC, normally closed; NO, normally open.

The airflow system is organised into three lines: a carrier line, a regulator line, and an odour line. The carrier line continuously delivers clean air to the output (*e.g.* a behavioural arena or face mask), providing stable baseline airflow. The regulator and odour lines share a common pressure regulator and are alternately routed via a 3-way solenoid valve. At rest, the airflow is directed thorough the regulator line; during odour delivery, airflow is redirected into the odour line and routed through the desired odour vials before merging with the carrier stream at the output.

Within the odour line, each module contains paired solvent and odour bottles that can be specifically delivered using individual solenoid valves. This modular architecture enables delivery of clean air, single odours, or defined mixtures of up to four odorants while maintaining stable output flow and pressure (Fig. 1). The design also allows straightforward expansion to additional odour modules if desired.

Logistically, we built the system on a small metal wheeled trolley (RA SKOG, IKEA), using it as a support platform. Its three wire mesh tiers facilitate stacking regulators and odour modules across the different levels, and the mesh allows tubing to pass through while keeping runs as short as possible. The wheeled design also allows for easy transport between experimental setups. Users should configure the platform and layout to suit their experimental requirements, with particular attention to minimising tubing length and maintaining a clear cable layout.

#### Air supply system

The olfactometer is supplied with compressed air from a medical air compressor (MGF compressors, S-OF100-003-MS2), although this may be substituted with an oxygen or medical air cylinder, or, if suitably clean, with pressure air from a laboratory supply.

From the compressor, the air is divided in two and enters two pressure regulators (SMC, AR20-F01BG-B) set to 0.1 MPa. One pressure regulator supplies the carrier line, while the second supplies both the regulator and odour lines (Fig. 1B).

The carrier line then passes through a dedicated flowmeter (Key Instruments, 25102A13BVBN) set to 1 L/min and flows directly and continuously to the output, reducing residual odour accumulation at the olfactometer’s output. The second pressure regulator then feeds a 3-way solenoid valve (Lee Company, LHDB1233418H) which splits the air into the regulator and odour lines. The regulator line is connected to the normally open output port of the 3-way valve and passes through a flowmeter set to 0.1 L/min and, when not delivering odour, flows with the carrier line directly to the output (Fig. 1B).

The odour line is connected to the normally closed output port of the 3-way valve. When odour delivery is triggered, the 3-way valve redirects airflow into the odour line, which distributes airflow through four parallel modules, each equipped with an individual flowmeter set to 0.1 L/min. Each module flowmeter is then connected to a speed controller (SMC, AS1211F-M5-04) and then to a normally closed 2-way solenoid valve (Lee Company, LHDB1262245D) positioned upstream of each of the solvent and odour bottles.

The exact flowmeter settings should be calibrated to match specific experimental conditions. For example, with the values aforementioned, at rest there will be 1.1L/min flowing to the output (1L/min from carrier line, 0.1L/min from regulator line). When odour is delivered there will still be 1.1L/min flowing to the output (1L/min from carrier line, 0.1L/min from odour line). However, if four odour bottles were to be delivered there would be an increase to 1.4L/min (1L/min from carrier line, 4 x 1L/min from the odour line). When delivering odour mixtures, the regulator line should be set to match the total flow expected from the odour line (*e.g.,* 0.4L/min when delivering odours from four bottles).

#### Odour modules

Our odour module design is adapted from (21). The system consists of four modules, each containing two sealed glass bottles: a solvent bottle and an odour bottle containing the diluted odorant. In total, the olfactometer accommodates eight bottles (four solvent and four odour), with modules colour-coded Red, Blue, Green, and Yellow for identification and reference (Fig. 1B). This system can be expanded by the introduction of any number of additional modules.

Each bottle is fitted with two needles: an inlet needle which provides air from the odour line and an outlet connected to the common final output (Fig. 1A). Individual bottles are engaged by normally closed solenoid valves positioned upstream of the inlet needles, allowing airflow to pass through a selected bottle where it becomes saturated with odour vapour before merging with the output stream.

A key feature of this design is the pairing of solvent and odour bottles within each module. During odour delivery, either the solvent bottle or the odour bottle of a given module should be opened, allowing the total number of active airflow channels (*i.e.,* open bottles) to remain constant across stimulus conditions. This approach ensures consistent airflow when presenting solvent controls, single odours, or odour mixtures, thereby avoiding pressure differences between conditions.

#### Electronic control

Odour delivery is controlled by solenoid valves actuated by an Arduino microcontroller (Arduino Mega 2560 R3), programmed via the Arduino integrated development environment (IDE, version 2.3.5) on a standard desktop computer (Fig. 1A). A valve is opened by setting the corresponding Arduino output pin to a logical “HIGH” state for the duration specified in the control code.

The four-module configuration described here uses eight 2-way solenoid valves (Lee Company, LHDB1262245D), in addition to one 3-way solenoid valve (Lee Company, LHDB1233418H) which controls regulator/odour line switching. The 2-way valves are operated using spike-and-hold circuits implemented on 2-way valve driver boards. The 3-way valve is operated via a dedicated 3-way valve driver board. Power for the spike-and-hold circuits is supplied by dedicated converter boards. Each converter board converts a 12 V input into two 7 V and two 2.8 V outputs, providing the voltages required for valve opening and holding currents. Two converter boards are sufficient to power the four-module system described here, although the design can be readily expanded to support additional odour modules if required.

Each 2-way valve driver board contains two identical circuits, allowing two solenoid valves (**i.e**., one module) to be operated from a single board. Consequently, four 2-way driver boards are required to control the eight 2-way valves in the Red, Blue, Green, and Yellow odour modules, while one separate driver circuit operates the 3-way valve (Fig. 2 and Supplementary Fig. 1).

**Fig 2.**
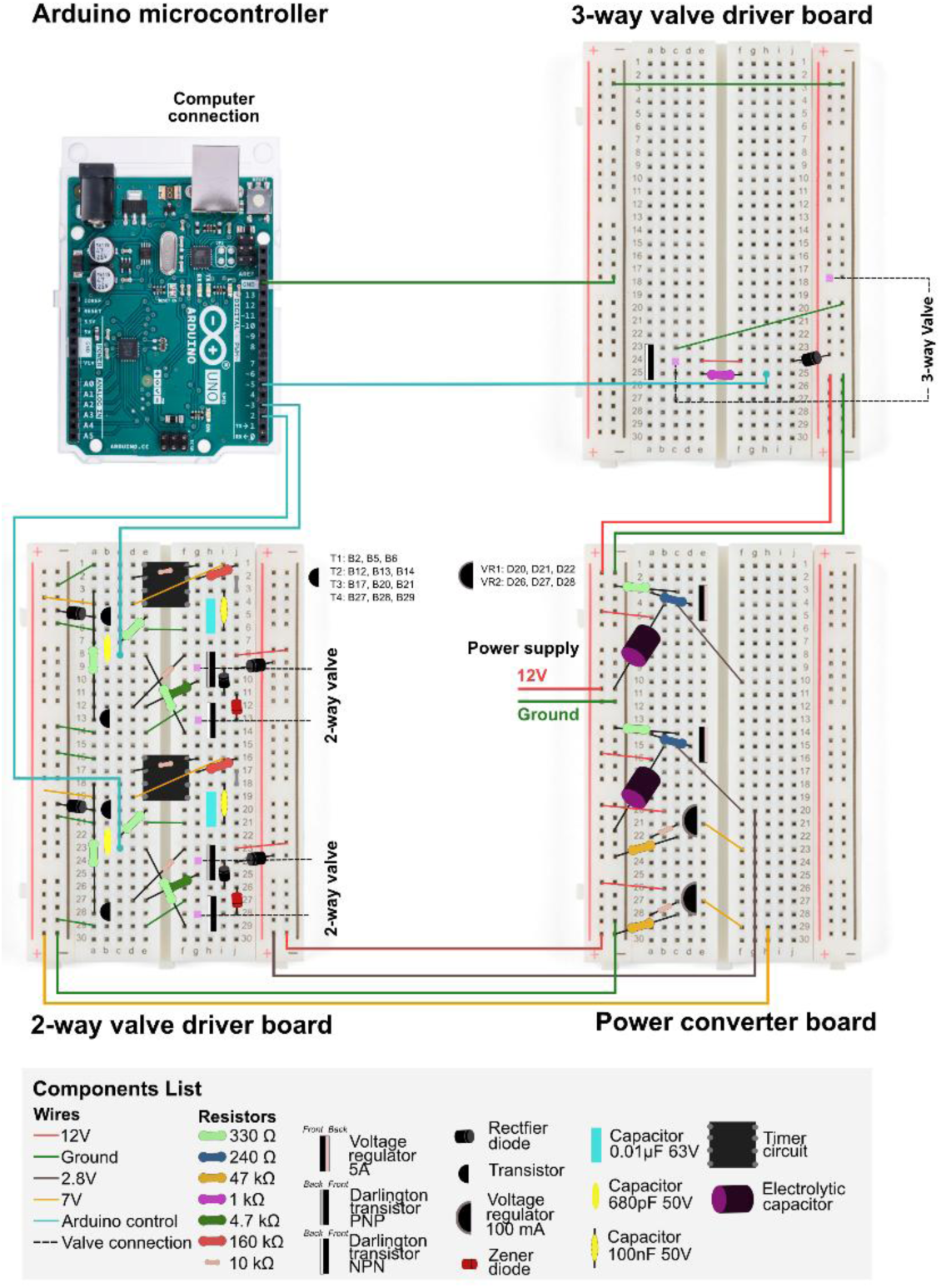
Breadboard layout and wiring for electronics used to control the olfactometer. **Arduino microcontroller and computer connection.** The Arduino provides digital output signals (blue wires) to control individual valves via the driver boards. **3-way valve driver board.** The circuit shown controls a single 3-way solenoid valve and is assembled on a dedicated breadboard. **2-way valve driver board**. Each 2-way valve driver board consists of two circuits: rows 1–14 control one valve, and rows 16–29 control a second valve. **Power converter board**. A 12 V input supply (red) is converted into regulated voltage lines that are distributed to the valve driver boards. Ground (green) is shared across all boards. Each converter board supplies power to two valve driver boards. A legend of all electronic components used (resistors, transistors, diodes, capacitors, and voltage regulators) is provided at the bottom of the figure. Further schematics of the circuits are presented in supplementary figure 1.

### Construction

#### Air supply system

See parts list in Table 1 and schematic in Fig. 3 for further guidance. Tubing lengths are provided here as a guide and should be optimised to the setup. Tubing should be kept as short and consistent across modules as possible to avoid low output or flow differences.

1. Take two pneumatic pressure regulators (SMC, AR20-F01BG-B). Attach G1/8-to-6 mm adaptors (RS Components, 299-3819) to the input ports and G1/8-to-4 mm adaptors (RS Components, 299-3796) to the output ports of each regulator (Fig. 3A, left).
2. Take a 6 mm Y-connector (RS Components, 916-0918) and attach two ~25 cm lengths of 6 mm diameter polyether tubing to the two-pronged end. Connect the other ends of these tubes to the input ports of the two pressure regulators (Fig. 3A, right). Using additional 6 mm tubing, connect the air compressor (MGF Compressors, S-OF100-003-MS2) to the remaining end of the Y-connector.
3. Take six flowmeters (Key Instruments, 25102A13BVBN) and securely attach two elbow adaptors (SMC, KQ2L03-34AS) to each flowmeter (Fig. 3Bi). One flowmeter will be used for the carrier line, one for the regulator line, and the remaining four for the odour modules. Colour-coding the odour flowmeters may help with identification (Fig. 3C).
4. Securely mount the six flowmeters to the shelf of a trolley or another suitable structure (Fig. 3Bii). Connect a ~30 cm length of 4 mm tubing from the output port of the carrier line pressure regulator to the bottom (input) elbow adaptor of the carrier-line flowmeter (Fig. 3C).
5. Connect a ~10 cm length of 4 mm diameter tubing to the output port of the regulator line and odour line pressure regulator. Using a 3/32” to 1/16” adaptor (Cole-Parmer, 6365-50), connect this tubing to a ~5 cm length of 2 mm diameter tubing. Attach the other end of the 2 mm tubing to the common (middle) input port of a 3-way solenoid valve (Lee Company, LHDB1233418H).
6. Connect two ~5 cm lengths of 2 mm tubing to the normally closed (NC, top) and normally open (NO, bottom) ports of the 3-way solenoid valve. Insert a 3/32” to 1/16” adaptor into the free end of each piece of tubing.
7. Connect a ~10 cm length of 4 mm tubing to the 3/32” to 1/16” adaptor attached to the NO port. Attach the other end of this tubing to the input elbow adaptor of the regulator line flowmeter.
8. Connect a ~5 cm length of 4 mm tubing to the 3/32” to 1/16” adaptor attached to the NC port. Form a 4mm-to-4mm double Y adaptor (by combining RS Components, 176-1294 with Festo, QSQ-G1/4-4) and attach the other end of this tubing to the single (non-branched) port of the double Y adaptor.
9. Using ~30 cm lengths of 4 mm tubing, connect each of the four odour-module flowmeters (via the bottom elbow adaptors) to the double Y adaptor (Fig. 3C).
10. Attach ~30 cm lengths of 4 mm tubing to the top (output) elbow adaptor of each of the four odour flowmeters. Connect each tube to a separate speed controller (SMC, AS1211F-M5-04).
11. Attach an M5-to-4 mm adaptor (RS Components, 176-1293) to each speed controller. Using a ~2 cm length of 4 mm tubing, connect each speed controller to a 4 mm Y connector (RS Components, 916-0915; Fig. 3Biii).
12. Attach ~20 cm lengths of 4 mm tubing to the two-pronged end of the 4 mm Y connector. These tubes will deliver air to the odour bottles (Fig. 3Biii).

**Fig 3.**
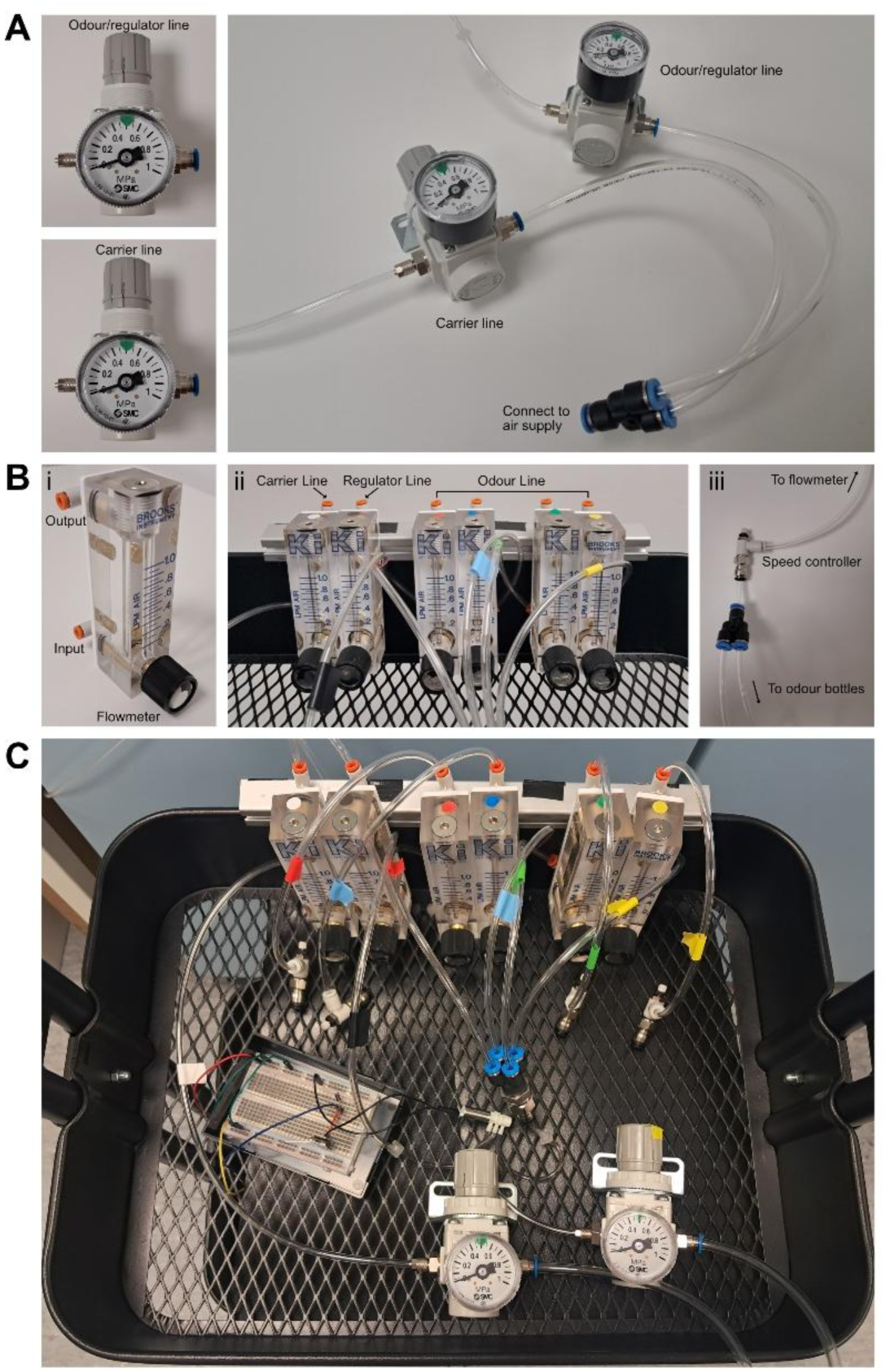
Air supply and regulation for the olfactometer. **(A)** Pressure regulators for the carrier and regulator/odour lines. Left: regulators fitted with input and output adaptors. Right: tubing connecting the regulators to the air supply (*e.g.,* an air compressor). **(B)** Flowmeter setup. i) Individual flowmeter with input and output connections indicated. ii) All six flowmeters mounted on the olfactometer trolley. iii) Speed controller connecting the flowmeter output to the odour bottles. **(C)** Complete air supply and regulation system, showing tubing connections between regulators, flowmeters, and speed controllers.

**Table 1 –.**
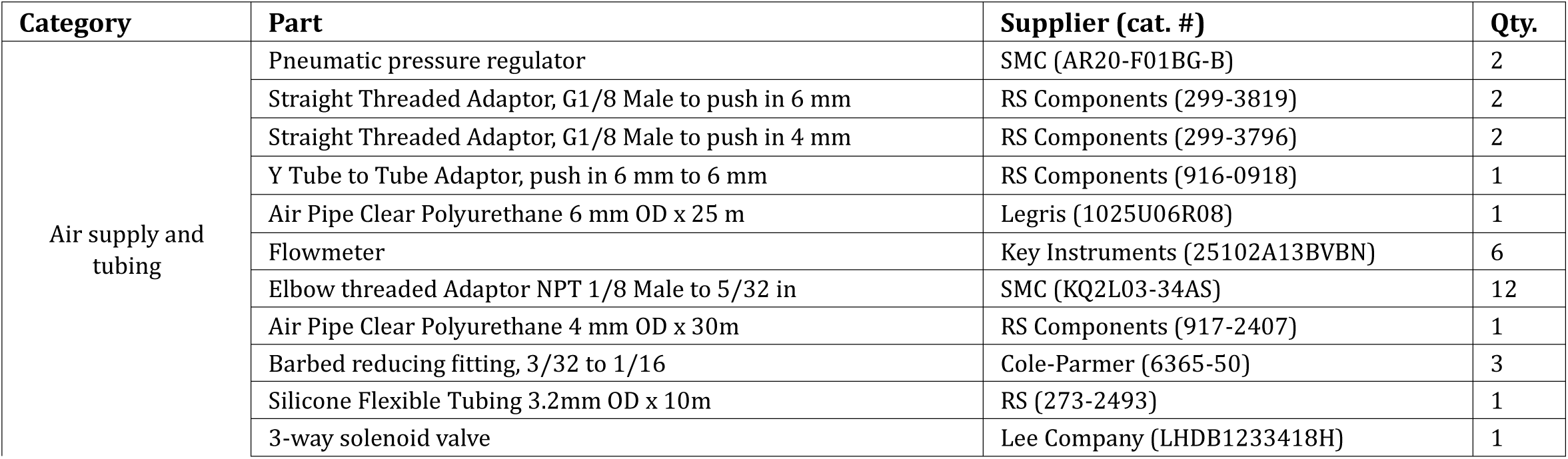
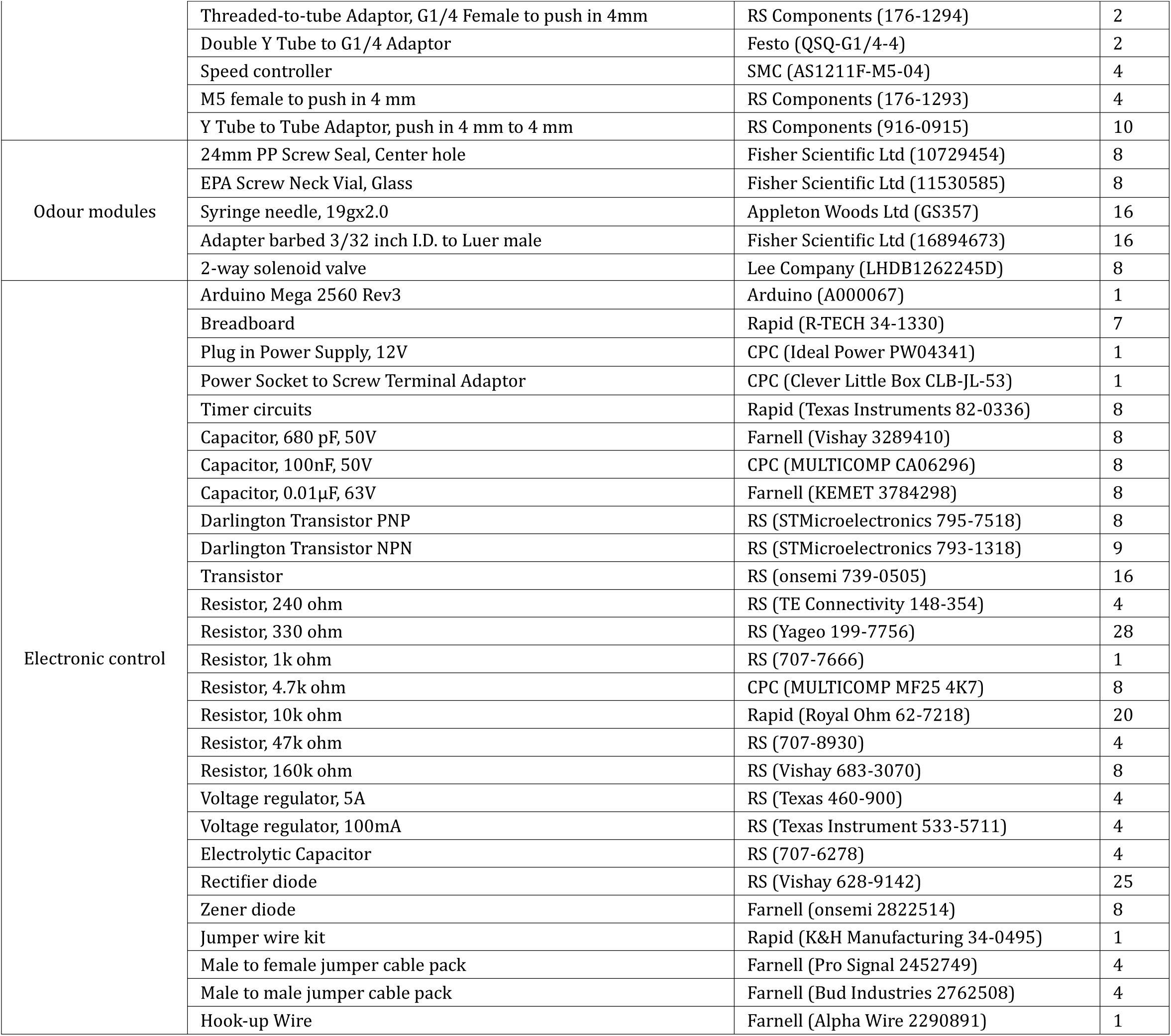
Olfactometer parts list. We include these product details for reference to aid replication of our design, not as endorsements of specific suppliers. Better or cheaper alternatives may be available elsewhere. Readers should research local suppliers and compare options when sourcing materials.

#### Odour modules

See parts list in Table 1 and schematic in Fig. 4 for further guidance. We selected these bottles, caps, and needles after testing multiple alternatives. Screw-on caps were essential as snap-on versions failed under operating pressure. The silicone-sealed caps maintain integrity when pierced while minimising leakage. The 40ml brown glass bottles provide optimal volume as larger bottles require excessive odorant for adequate vapor concentration, while smaller ones cannot withstand operating pressures without valve failure nor enable the use of solid material (*e.g.*, coffee beans). Brown glass protects light-sensitive odorants. While our testing was thorough, other suitable options may exist.

1. Attach screw-top lids (Fisher Scientific Ltd, 10729454) to eight odour bottles (Fisher Scientific Ltd, 11530585).
2. Carefully insert two 19G syringe needles (Appleton Woods Ltd, GS357) through the lid of each bottle (Fig. 4, top left). Remove the lid with the needles attached and clear any rubber debris from the needles by blowing air through.
3. Attach a barbed 3/32”–to–male Luer adaptor (Fisher Scientific Ltd, 16894673) to the inlet of each needle (Fig. 4, top left).
4. Designate one needle on each bottle as the *inlet* and the other as the *outlet*. Attach a short (~2 cm) length of 4 mm tubing to the outlet needle. This tubing will carry odourised air from the bottle (Fig. 4, middle).
5. Connect the tubing from the speed controller (Air flow control, step 11) to the input port of a solenoid valve (Lee Company, LHDB1262245D; Fig. 4, bottom left).
6. Attach a ~10 cm length of 4 mm tubing to the output port of the solenoid valve and connect the other end to the inlet needle of the odour bottle. This tubing delivers airflow into the bottle when the valve is activated.
7. Attach a ~15 cm length of 4 mm tubing to the outlet needle of the bottle. This tubing forms the module output line (Fig. 4, right).
8. Use a 4 mm Y connector (RS Components, 916-0915) to combine the output tubing from one solvent bottle and one odour bottle, forming a single odour module (Fig. 4, right).
9. Repeat steps 4–8 to assemble the remaining three odour modules.
10. Use a double-Y tube-to-tube adaptor (RS Components, 176-1294 and Festo, QSQ-G1/4-4) to combine the four module output lines into a single odour line.
11. Use a 4 mm Y connector to join the combined odour line with the top elbow adaptor from the carrier-line flowmeter.
12. Attach a length of 4 mm tubing to the remaining port of this Y connector. This tubing delivers the final odour mixture to the behavioural arena.
13. Before starting the experiment, add solvent to the solvent bottles and odour diluted in solvent to the odour bottles. We typically use 1 ml per bottle, although this can be adjusted provided the liquid level remains below the needle tips. Solid odour sources (e.g., coffee beans) may also be used.

**Fig 4.**
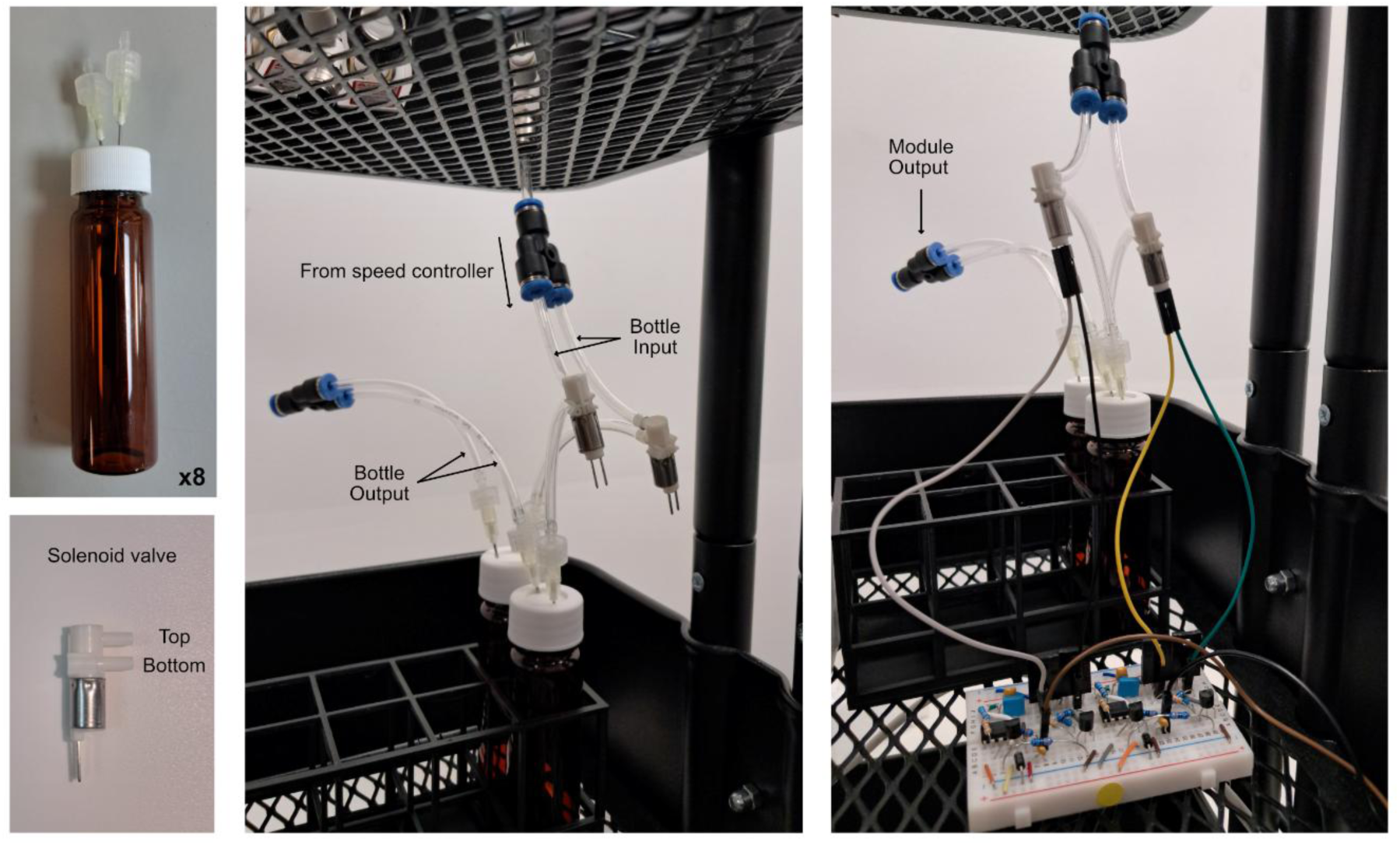
Odour and solvent bottle assembly. Top left: capped odour bottle with attached needles. Bottom left: Valve with connections labelled. Middle: odour and solvent bottles for the Red module, with tubing from Fig 3Biii connected to the bottle inputs. Right: connection between bottles and solenoid valves mounted on the valve driver board.

#### Electronic control

1. Assemble two power converter boards using the components in Table 1 and the schematic in Fig. 2. No soldering is required; insert the pins into the appropriate sockets.
2. Assemble one 3-way valve driver board and four 2-way valve driver boards using the same components list and schematic (Fig. 2).
3. Connect the power converter boards to the valve driver boards using jumper wires as shown in Fig. 2.
4. Connect the power supply to one power converter board (Fig. 2), then link this board to the second converter board so that both receive 12 V and GND.
5. Connect the Arduino digital output pins to the valve driver boards (Fig. 2) and connect Arduino GND to the GND rail of a power converter board.
6. Connect the Arduino (Arduino Mega 2560 R3) to a desktop PC or laptop via USB.

A construction troubleshooting mini-guide is provided in the supplementary material (S2).

### Software for performing experiments

#### Controlling odour delivery

1. Download and install the Arduino IDE (https://arduino.cc/en/software). Open the IDE and select the correct board model under *Tools > Board*.
2. Write a custom script defining the desired odour delivery sequence. Ensure that the digital pin assignments in the code correspond to the physical wiring of the valve driver boards.
3. Switch on the 12 V power supply. Upload the script to the Arduino using the *Upload* function in the Arduino IDE to begin odour delivery.

Example Arduino code for controlling the solenoid valves is provided below. For additional information on Arduino programming, see https://docs.arduino.cc/software/ide/.

int masterPin = 2; // Defines digital pin 2 as the pin used to control the 3-way valve

int odourPin = 3; // Defines digital pin 3 as the pin used to control the 2-way valve

void setup() {

pinMode(masterPin, OUTPUT); // Declares pin 2 an output

pinMode(odourPin, OUTPUT); // Declares pin 3 an output

digitalWrite(masterPin, LOW); // Air flows through NO output

digitalWrite(odourPin, LOW); // Valve initially closed

}

void loop(){

digitalWrite(masterPin, HIGH); // Diverts air through NC output

digitalWrite(odourPin, HIGH); // Opens odour valve

delay(1000); // Valves open for 1s

digitalWrite(masterPin, LOW); // Air returns to NO output

digitalWrite(odourPin, LOW); // Valve closed

delay(10000); // Wait 10s before next cycle

}

### Technical validation of olfactometer performance

#### Reliable odour delivery across repeated trials

We first validated whether our olfactometer could deliver reliable odour signals that were both stable across repeated presentations and tightly coupled to valve actuation using a miniPID (Aurora Scientific, 200C). A PID (photoionization detector) is a fast-response sensor that uses UV light to ionize volatile organic compounds and measure their concentration in real time; the resulting ionization generates a small current that is converted into a voltage signal (22). Importantly, although specific odours can give rise to characteristic waveform signatures on the PID signal,, PIDs do not identify which odour is present, only how much is there and for how long.

We recorded mini responses to ten consecutive 10s presentations of 1ml 5% methyl tiglate (MT) in diethyl phthalate solvent (sol). The traces exhibited a square-shaped waveform, with a rapid increase in signal following valve opening, a stable signal during odour delivery, followed by a return to baseline upon valve closure (Fig. 5A). Quantification (Fig. 5Bi-iv) confirmed consistent latency to odour onset (mean ± SD, 252.6 ± 13.1ms), peak amplitude (mean ± SD, 47.47 ± 3.22mV), area under the curve (mean ± SD, 4439641 ± 227436mV·ms), and decay time (mean ± SD, 1.62 ± 0.03s) across the 10 odour presentations. The greatest variability in signal was during the initial valve opening, which likely stems from this being the first time air enters the odour line, pressurising it, as discussed in (16).

**Fig 5.**
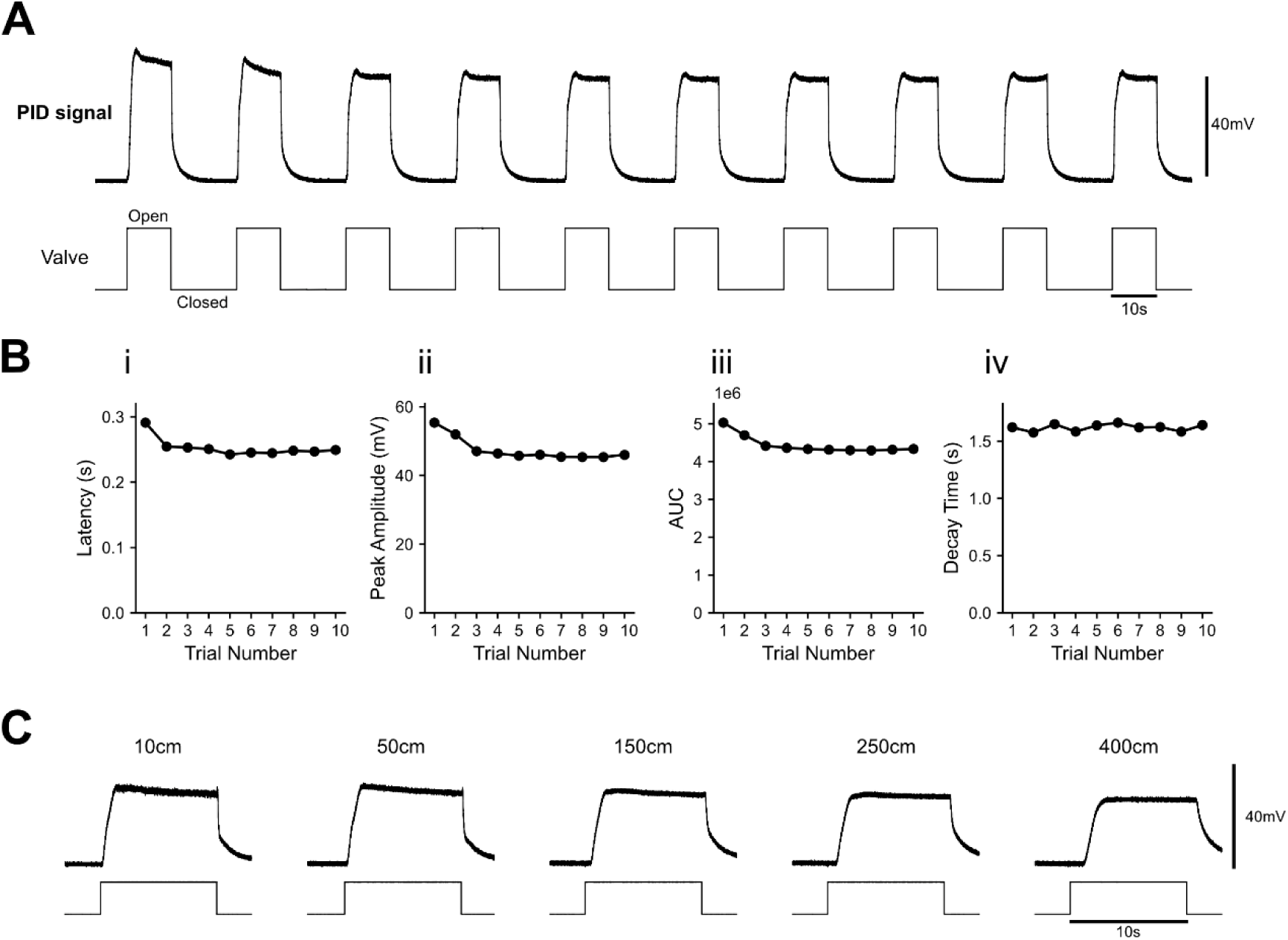
Odour delivery controlled by the olfactometer. **(A)** Complete recording trace of 5% methyl tiglate (MT) from a single module across ten 10s presentations. Top: miniPID signal; bottom: digital output voltage trace from Arduino indicating the timing of valve opening. **(Bi–iv)** Quantified features include (i) latency to odour onset, (ii) peak amplitude, (iii) area under the curve during valve opening, and (iv) decay time. **(C)** Example miniPID traces during 10s exposure of 5% MT across different output tubing lengths. Top: miniPID signal; bottom: digital output voltage trace from Arduino indicating the timing of valve opening. Tubing length is indicated above each trace. Full quantification of tubing length effect on odour latency in supplementary figure 3.

Since a key determinant of miniPID signal is the distance of the sensor from the odour (18–21,23,24), we sought to test whether the detected signal varies depending on the length of the tubing used prior to the miniPID sensor. We used tubing lengths ranging from 10cm to 4m in an attempt to capture what responses might look like across various set ups, including delivering stimuli into an fMRI where the olfactometer body has to be placed in a separate room due to metal components. Fig. 5C shows example traces from a range of tubing lengths. Quantification (Supplementary Fig. 3) reveals that while latency to odour onset and decay time are impacted by tubing length, there is minimal effect of tubing length on the actual amount of odour being delivered. Moreover, odour onset delay scales linearly with tubing length (R² = 0.917), allowing reliable predictions of latency for customized lengths.

In summary, the olfactometer reliably generates detectable odour signals with short latencies following valve actuation and high trial-to-trial stability.

#### Precise control of odour delivery duration

A key requirement of olfactometer performance is the ability to precisely control the duration of odour delivery. As such, we next sought to validate that our custom-olfactometer can reliably deliver odours across a range of stimulus presentation durations.

Using 5% MT delivered from one odour module, we recorded miniPID responses to ten consecutive presentations at valve opening durations ranging from 100ms to 1000ms. Reliable responses could be detected with opening times as low as 250ms (Fig. 6A), given our current design and flow settings. Across all opening times that produced reliable signals, there was minimal difference in odour onset latency (Fig 6Bi; 250ms = 283.9 ± 30.2ms, 500ms = 257.1 ± 11.3ms, 750ms = 248.8 ± 11.0ms, 1000ms = 250.5 ± 15.8ms). As opening duration increased there was a consistent increase in peak amplitude (250ms = 5.81 ± 0.60mV, 500ms = 23.3 ± 0.66mV, 750ms = 32.4 ± 0.40mV, 1000ms = 36.3 ± 0.33mV), area under the curve (250ms = 56577 ± 4215mV·ms, 500ms = 194432 ± 8179mV·ms, 750ms = 337169 ± 5429mV·ms, 1000ms = 439154 ± 5411mV·ms) and decay time (250ms = 0.37 ± 0.06s, 500ms = 0.79 ± 0.05s, 750ms = 1.24 ± 0.08s, 1000ms = 1.32 ± 0.05s).

**Fig 6.**
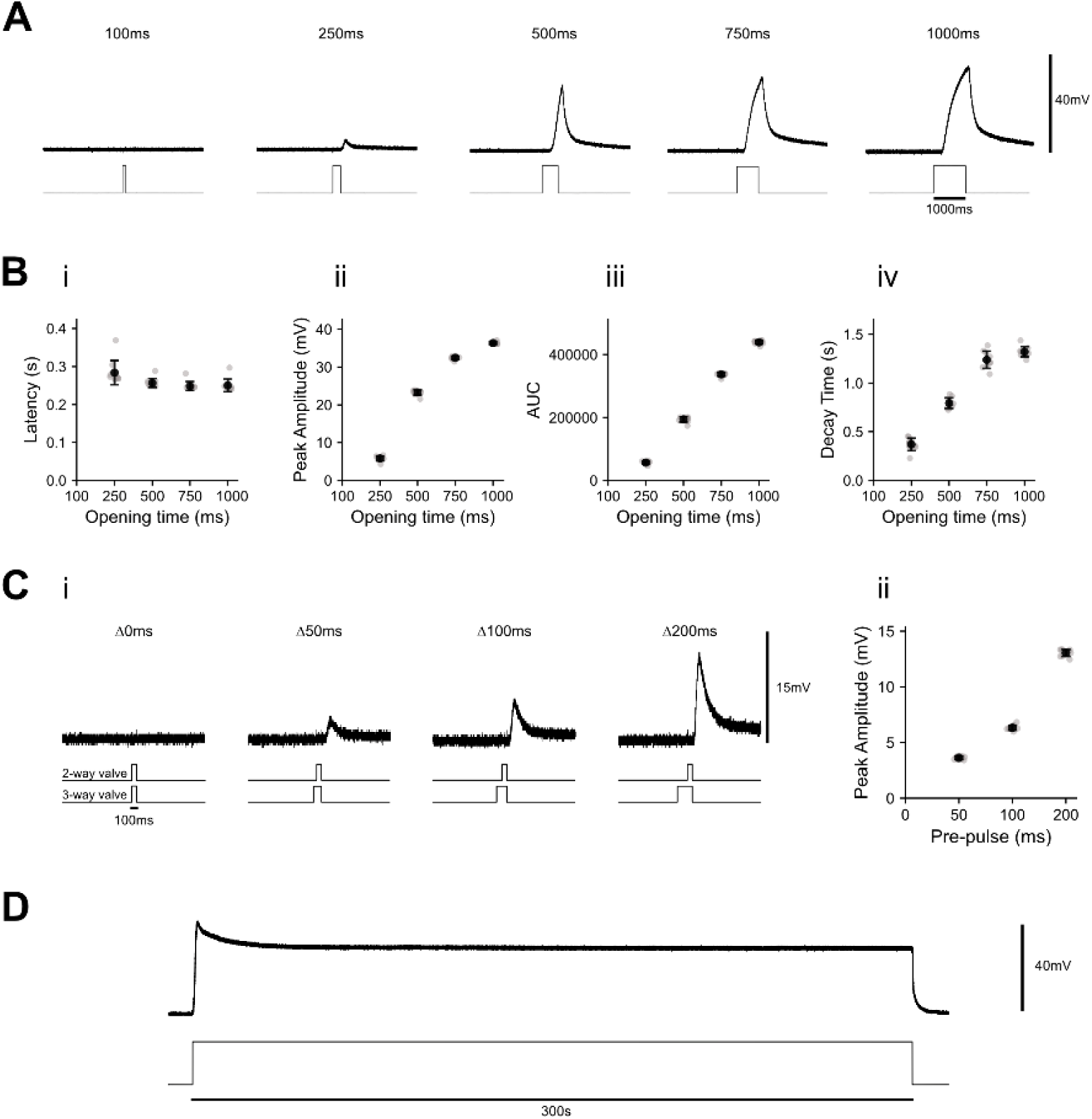
Odour delivery across different valve opening times. **(A)** Representative traces of 5% MT during presentations of increasing duration. Top: miniPID signal; bottom: digital output voltage trace indicating valve state. Valve opening duration is indicated above each trace. **(B)** Quantified features include (i) latency to odour onset, (ii) peak amplitude, (iii) area under the curve during valve opening, and (iv) decay time. **(C)** (i) Representative traces of 5% MT during 100ms presentations while varying pre-pulse duration. Top: miniPID signal; bottom: digital output voltage trace indicating valve state. Pre-pulse duration is indicated above each trace. (ii) Quantification of peak amplitude. **(D)** Trace of 5% MT during a 5-min continuous valve opening. Top: miniPID signal; bottom: digital output voltage trace. Grey circles are individual repetitions, black circles and lines are mean ± SD.

In the recordings described above, the 3-way valve (which directs airflow through either the regulator or odour line) and the 2-way solenoid valve connected to the MT odour bottle were actuated simultaneously. The absence of a detectable signal in the 100ms condition may reflect insufficient time for airflow to propagate from the 3-way valve junction into the odour bottle and to the output. To test this, we introduced an offset between activation of the 3-way valve and the odour valve, such that, the 3-way valve was opened prior to the odour valve (pre-pulse), allowing airflow to establish within the odour line before odour release. We tested pre-pulse durations of 50ms, 100ms, and 200ms, in addition to the 100ms condition with simultaneous valve activation (0 ms pre-pulse). As seen above, no reliable signal was detected when the 3-way and 2-way valves were actuated simultaneously (0ms pre-pulse). In contrast, introducing a pre-pulse produced reliable odour signals, with peak amplitude increasing systematically with pre-pulse duration (Fig. 6C; 50ms pre-pulse = 3.64 ± 0.13mV, 100ms pre-pulse = 6.33 ± 0.21mV, 200ms pre-pulse = 13.05 ± 0.29mV at 200ms). These results indicate that a brief pre-pulse enhances odour output at short presentation durations by allowing airflow to reach the odour bottle prior to release.

Having established reliable odour delivery at short durations, we next assessed performance during longer stimulation periods commonly used in both human and animal behavioural experiments(25,26). Specifically, we sought to determine whether the small liquid volume used in each odour vial (1ml) was sufficient to sustain consistent output without significant signal depletion during prolonged odour delivery.

To test this, we recorded miniPID responses during a continuous 5-min presentation of 5% MT (Fig. 6D). Following an initial transient peak associated with valve opening and odour onset, the signal rapidly stabilised and reached a sustained plateau. This plateau remained highly stable throughout the remainder of the stimulation period, with negligible decline in signal amplitude. These results indicate that the odour volume used is sufficient to support prolonged odour delivery without appreciable depletion, confirming the suitability of the system for behavioural paradigms requiring extended stimulus presentations.

#### Controlled titration of odour concentration

In addition to precise temporal control, effective olfactometers must provide reliable control over odour concentration. While this is traditionally achieved by placing the same odour in different bottles at varying liquid dilutions, in our system output concentration can instead be modulated by adjusting the relative flow rates of the carrier and odour lines, thereby varying the degree of air dilution while keeping the liquid odour concentration constant. We therefore evaluated whether systematic changes in the carrier-to-odour flow ratio produce predictable and stable changes in odour signal amplitude.

To test this, we systematically varied the carrier-to-odour flow ratio while maintaining a constant total flow rate of 3L/min. Carrier flow ranged from 2.9L/min with 0.1L/min odour flow (29:1 ratio) to 1.5L/min carrier with 1.5L/min odour flow (1:1 ratio). The odour bottle contained 5% MT, and miniPID responses were recorded during ten consecutive 10s presentations at each dilution ratio. Systematic variation of the carrier-to-odour flow ratio produced predictable changes in miniPID signal characteristics, consistent with graded control of odour concentration (Fig. 7A). As the proportion of odour flow increased, peak amplitude increased monotonically (Fig. 7B; 29:1 = 13.71 ± 0.72mV, 9:1 = 20.49 ± 1.02mV, 5:1 = 25.03 ± 1.70mV, 2:1 = 31.42 ± 2.82mV, 1:1 = 37.44 ± 3.12mV). Similarly, area under the curve increased with increasing odour flow (29:1 = 1.18 × 10^6^ ± 3.51 × 10^4^mV·ms, 9:1 = 1.76 × 10^6^ ± 3.52 × 10^4^mV·ms, 5:1 = 2.15 × 10^6^ ± 5.99 × 10^4^, 2:1 = 2.70 × 10^6^ ± 9.19 × 10^4^mV·ms, 1:1 = 3.24 × 10^6^ ± 9.72 × 10^4^mV·ms). Odour onset latency decreased as odour flow increased (29:1 = 0.385 ± 0.129s, 9:1 = 0.295 ± 0.070s, 5:1 = 0.226 ± 0.050s, 2:1 = 0.180 ± 0.022s, 1:1 = 0.148 ± 0.015s), while decay time increased (29:1 = 0.590 ± 0.065s, 9:1 = 0.714 ± 0.062s, 5:1 = 0.718 ± 0.034s, 2:1 = 0.744 ± 0.038s, 1:1 = 0.832 ± 0.036s). Together, these results demonstrate that adjusting the carrier-to-odour flow ratio provides stable and graded control over odour concentration at the output.

**Fig 7.**
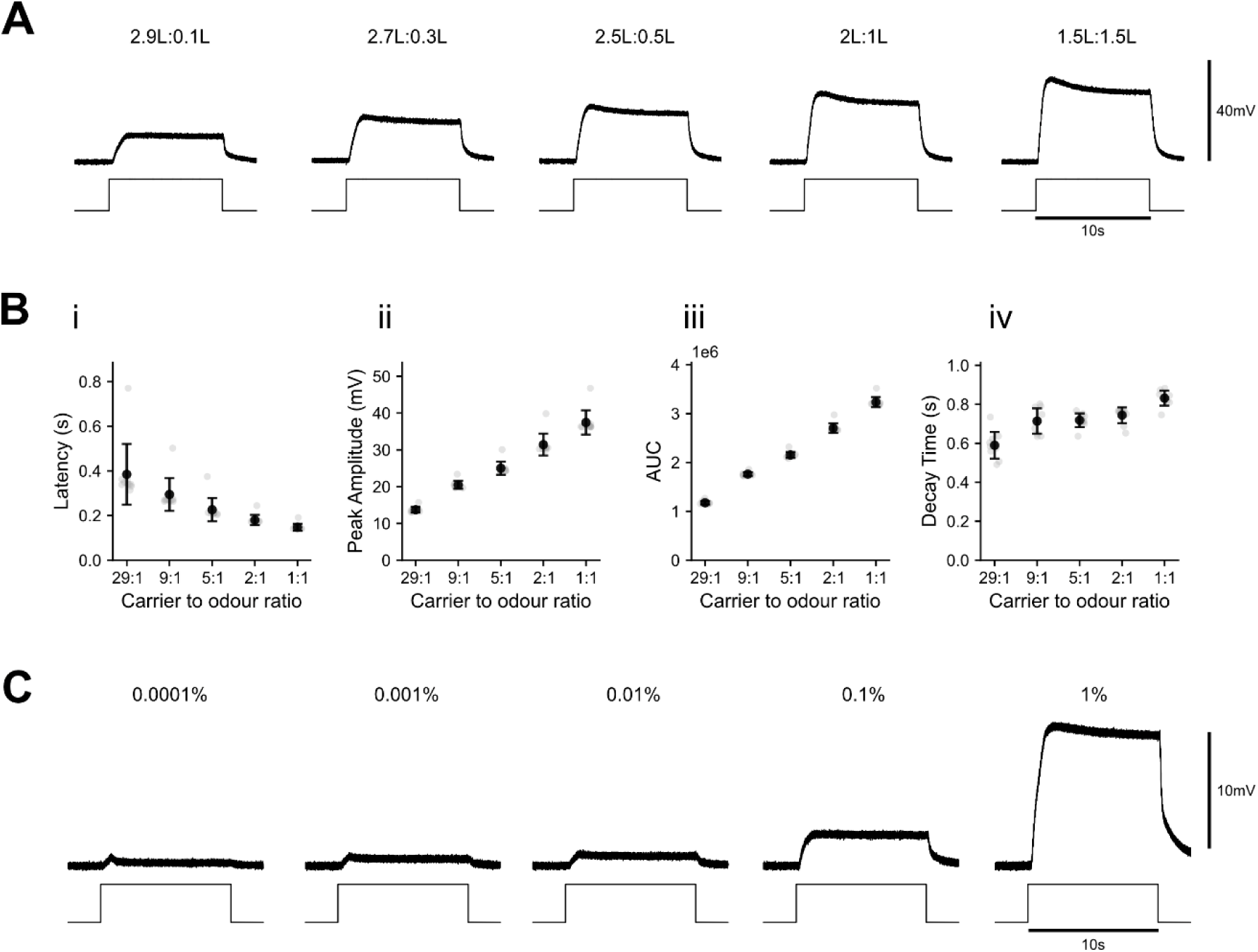
Odour delivery across different air and liquid dilutions. **(A)** Representative traces of 5% MT during presentations at differing carrier-to-odour flow ratios. Top: miniPID signal; bottom: digital output voltage trace indicating valve state. Top: miniPID signal; bottom: digital output voltage trace indicating valve state. Carrier-to-odour flow ratio is indicated above each trace. **(B)** Quantified features include (i) latency to odour onset, (ii) peak amplitude, (iii) area under the curve during valve opening, and (iv) decay time. Grey circles are individual repetitions, black circles and lines are mean ± SD. **(C)** Averaged MiniPID responses to methyl tiglate at different liquid dilutions. Traces show the mean of 20 presentations per concentration. Top: miniPID signal; bottom: digital output voltage trace. Odour liquid dilution is indicated above each trace.

In addition to air dilution, odour concentration can also be more traditionally controlled by varying the liquid-phase dilution of the odorant within the odour bottle. To assess the sensitivity and lower detection limits of the system, we tested a range of MT dilutions (0.0001%, 0.001%, 0.01%, 0.1%, and 1%). MiniPID responses were recorded during repeated 10s odour presentations at each concentration. At lower concentrations, individual presentations produced signals that were close to the noise floor of the detector and were not reliably distinguishable on a trial-by-trial basis. To improve signal detection, each concentration was presented 20 times and the resulting traces were averaged. Averaging revealed progressively larger and more distinct signals observed as liquid concentration increased (Fig. 7C). These results confirm that odour concentration can be effectively controlled through liquid dilution, complementing the air dilution approach described above.

#### Consistent delivery across the four odour modules

Our olfactometer consists of four independent modules, allowing the delivery of up to four distinct odours within a single experiment (although this is expandable). The different modules can be activated sequentially, as, for example, in a habituation-dishabituation task where a subject is exposed repeatedly to one odour before being exposed to a new odour (27). Having demonstrated reliable repeatability within one modules (red module, Fig. 5), we next test inter-module consistency by delivering the same odour from multiple modules. To do this, we delivered 5% MT from each of the four modules (10s odour presentation, 15s intertrial interval, repeated 10 times per module). The four modules showed similar waveforms, indicating similar odour presentation dynamics across the four modules (Fig. 8A).

**Fig 8.**
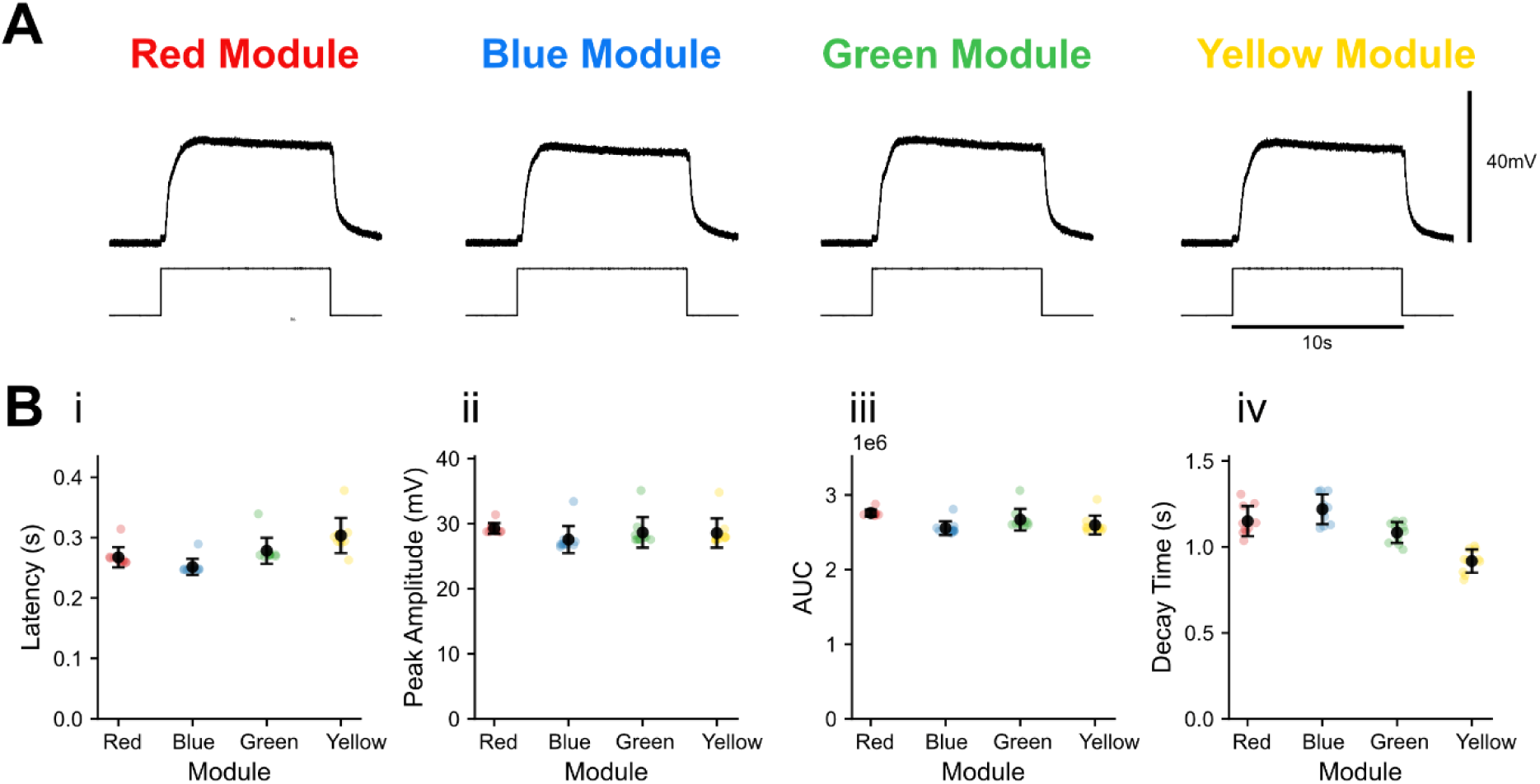
Odour delivery across different odour modules. **(A)** Representative traces of 5% MT from all four modules during 10s presentations with a 15s inter-stimulus interval. Top: miniPID signal; bottom: digital output voltage trace indicating valve state. **(B)** Quantified features include (i) latency to odour onset, (ii) peak amplitude, (iii) area under the curve during valve opening, and (iv) decay time. Black circles and lines are mean ± SD, coloured circles are individual repetitions. Recording traces of sequential 30s presentations of 5% MT from all four modules are presented in supplementary figure 4.

As observed previously, trial-to-trial variability was low across all modules, indicating that each module is capable of reliable and reproducible odour delivery (Fig. 8Bi–iv). Peak amplitudes were highly consistent between modules (Red = 29.26 ± 0.78mV, Blue = 27.55 ± 1.97mV, Green = 28.64 ± 2.22mV = Yellow: 28.56 ± 2.14mV). Similarly, area under the curve values were comparable across modules (Red: 2.76 × 10^6^ ± 4.23 × 10^4^mV·ms, Blue: 2.56 × 10^6^ ± 8.78 × 10^4^mV·ms, Green: 2.67 × 10^6^ ± 1.37 × 10^5^mV·ms, Yellow: 2.60 × 10^6^ ± 1.20 × 10^5^mV·ms), demonstrating consistent overall odour delivery. Odour onset latency was also similar between modules (Red = 0.267 ± 0.016s, Blue = 0.252 ± 0.013s, Green = 0.278 ± 0.020s, Yellow = 0.304 ± 0.028s), indicating comparable delivery timing. Decay times showed modest variation but remained within a narrow range (Red = 1.150 ± 0.083s, Blue = 1.219 ± 0.082s, Green = 1.084 ± 0.058s, Yellow = 0.917 ± 0.064s), consistent with stable and repeatable odour clearance dynamics. These results confirm that odour delivery is highly consistent across modules, demonstrating the reliability and potential scalability of the multi-module olfactometer design.

#### Generation of odour mixtures

Our olfactometer’s modular architecture enables both sequential and simultaneous activation of multiple modules, allowing delivery of defined odour mixtures for studies of odour de-mixing and signal–background segregation (21,28–30). Unlike systems that require physical mixing of liquid odorants prior to vaporisation (31,32), mixtures are generated by selectively opening individual odour bottles, allowing vapours to combine within the airflow downstream of the modules. This approach enables flexible and temporally precise control over mixture composition. However, because each open bottle contributes airflow to the output, activating different numbers of bottles may alter the total pressure at the output (Fig. 9A). This is an important point to consider, as olfactory sensory neurons as well as downstream neurons in the olfactory bulb are sensitive to mechanical stimulation (33,34), and unintended pressure differences may confound behavioural or physiological measurements. To quantify this effect, we measured pressure and miniPID responses during delivery of MT alone and in combination with one, two, or three solvent channels.

**Fig 9.**
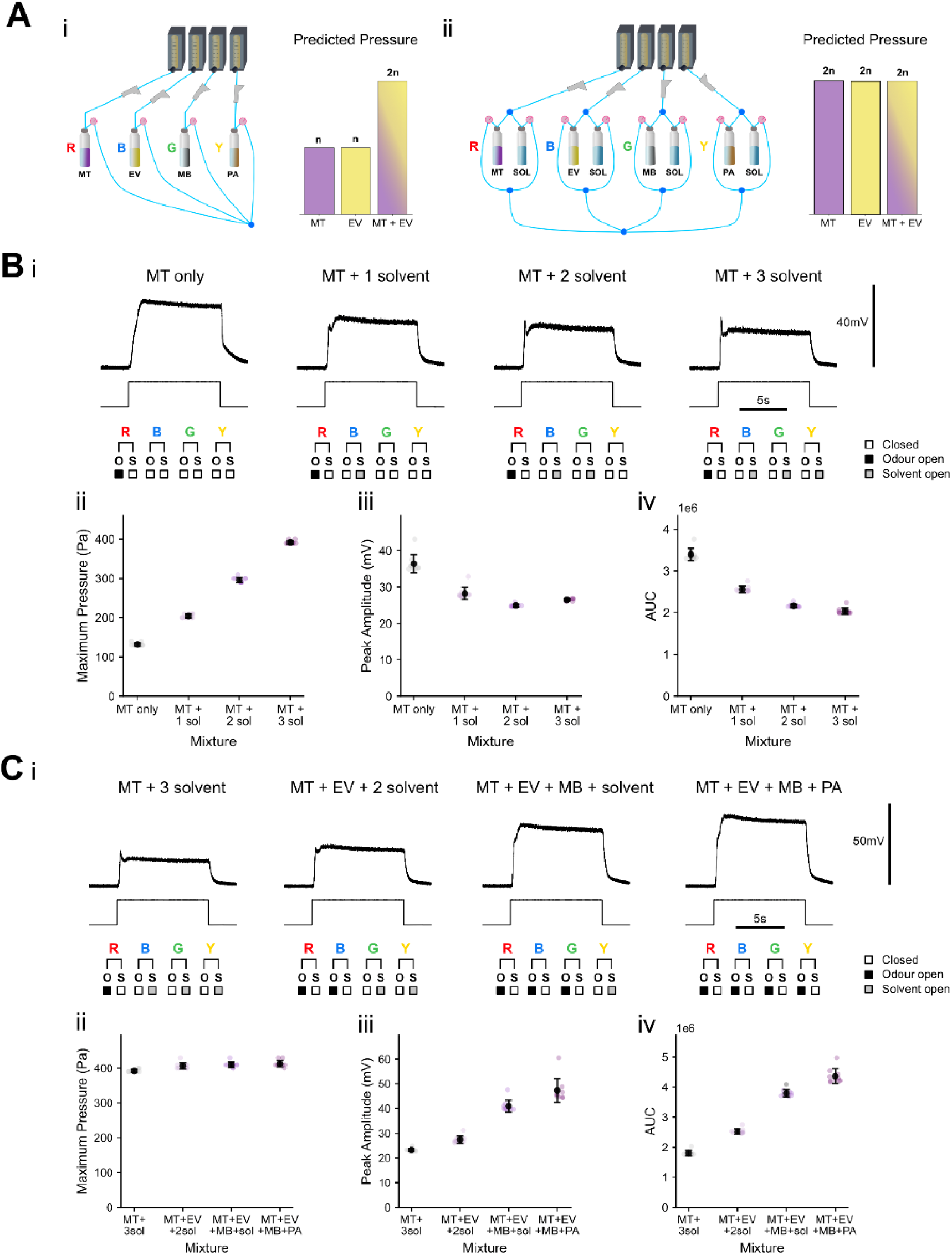
Delivery of odour mixtures. **(Ai)** Schematic of an olfactometer lacking solvent channels and the corresponding predicted pressure during mixture delivery. **(Aii)** Our olfactometer schematic and corresponding predicted pressure during mixture delivery. **(Bi)** Representative traces of 5% MT while increasing the number of co-activated solvent bottles. Top: miniPID signal; middle: digital output voltage trace indicating valve state; bottom: schematic showing which modules and bottles are active for each condition. (Bii–iv) Quantified features include (i) maximum pressure, (ii) peak amplitude, and (iii) area under the curve. Different timing of delivery are presented in supplementary figure 5. **(Ci)** Representative traces of different odour mixtures. Top: miniPID signal; middle: digital output voltage trace indicating valve state; bottom: schematic showing which modules and bottles are active for each condition. Mixture composition is indicated above each trace. (Cii–iv) Quantified features include (i) maximum pressure, (ii) peak amplitude, and (iii) area under the curve. MT, methyl tiglate; EV, ethyl valerate; MB, methyl butyrate; PA, pentyl acetate; SOL, solvent. Ligh purple circles are individual repetitions, black circles and lines are mean ± SD.

Peak pressure increased systematically with the number of open bottles (Fig. 9B; MT only = 132 ± 4Pa, MT + 1 solvent = 204 ± 4.9Pa, MT + 2 solvent = 296 ± 6.6Pa, MT + 3 solvent = 392 ± 4Pa), confirming that output pressure is directly determined by the number of active channels. Despite this increase in airflow, odour responses remained robust under all conditions. Peak amplitude decreased with the addition of solvent channels (MT only = 36.41 ± 2.36mV, MT + 1 solvent = 28.23 ± 1.59mV, MT + 2 solvent = 24.90 ± 0.35mV, MT + 3 solvent = 26.46 ± 0.26mV), and area under the curve showed a similar trend (MT only = 3.39 × 10^6^ ± 1.35 × 10^5^, MT + 1 solvent = 2.56 × 10^6^ ± 7.08 × 10^4^, MT + 2 solvent = 2.16 × 10^6^ ± 3.97 × 10^4^, MT + 3 solvent = 2.04 × 10^6^ ± 7.31 × 10^4^). Importantly, clear odour signals were detectable even when three solvent channels were co-activated.

To stabilise pressure across experimental conditions, the olfactometer supports solvent balancing, implemented through the Arduino control software, whereby modules not delivering odour instead deliver solvent. This ensures that a constant number of bottles remain open regardless of mixture composition. To verify that this approach effectively stabilises pressure, we compared responses across mixtures of increasing complexity while maintaining four active channels in all conditions (Fig. 9C). Pressure remained constant across conditions (MT + 3 solvent = 392 ± 4Pa, MT + EV + 2 solvent = 407 ± 9Pa, MT + EV + MB + solvent = 410 ± 7.7Pa, MT + EV + MB + PA = 413 ± 9Pa), confirming that pressure is determined by the number of open channels rather than odour identity. In contrast, both peak amplitude (MT + solvent = 23.20 ± 0.61mV, MT + EV + 2 solvent = 27.38 ± 1.39mV, MT + EV + MB + solvent = 40.94 ± 2.29mV, MT + EV + MB + PA = 47.29 ± 4.59mV) and area under the curve (MT + 3 solvent = 1.80 × 10^6^ ± 8.09 × 10^4^mV·ms, MT + EV + 2 solvent = 2.52 × 10^6^ ± 8.38 × 10^4^mV·ms, MT + EV + MB + solvent = 3.80 × 10^6^ ± 1.13 × 10^5^mV·ms, MT + EV + MB + PA = 4.36 × 10^6^ ± 2.31 × 10^5^mV·ms increased with the number of odour channels, as expected from the greater total odour delivered.

To stabilise pressure across experimental conditions, the olfactometer supports solvent balancing, implemented through the Arduino control software, whereby modules not delivering odour instead deliver solvent. This ensures that a constant number of bottles remain open regardless of mixture composition. To verify that this approach effectively stabilises pressure, we compared responses across mixtures of increasing complexity while maintaining four active channels in all conditions (Fig. 9C). Pressure remained constant across conditions (MT + 3 solvent = 392 ± 4Pa, MT + EV + 2 solvent = 407 ± 9Pa, MT + EV + MB + solvent = 410 ± 7.7Pa, MT + EV + MB + PA = 413 ± 9Pa), confirming that pressure is determined by the number of open channels rather than odour identity. In contrast, both peak amplitude (MT + solvent = 23.20 ± 0.61mV, MT + EV + 2 solvent = 27.38 ± 1.39mV, MT + EV + MB + solvent = 40.94 ± 2.29mV, MT + EV + MB + PA = 47.29 ± 4.59mV) and area under the curve (MT + 3 solvent = 1.80 × 10^6^ ± 8.09 × 10^4^mV·ms, MT + EV + 2 solvent = 2.52 × 10^6^ ± 8.38 × 10^4^mV·ms, MT + EV + MB + solvent = 3.80 × 10^6^ ± 1.13 × 10^5^mV·ms, MT + EV + MB + PA = 4.36 × 10^6^ ± 2.31 × 10^5^mV·ms increased with the number of odour channels, as expected from the greater total odour delivered.

Together, these results demonstrate that output pressure is governed by the number of active modules, whereas odour signal magnitude is determined by odour composition. The ability to maintain a constant number of open channels using solvent balancing therefore ensures stable pressure while preserving precise and flexible control over odour mixtures, thereby validating the modular design of our olfactometer.

### Biological validation

#### Mouse behaviour

Having validated the performance of our olfactometer with extensive PID recordings, we next investigated the suitability of our olfactometer for mouse behavioural testing. To do this, we utilised a habituation-dishabituation task, a standard task used to assess the ability of a mouse to discriminate between two odours (27,35). Mice were placed in a custom-built rectangular arena made of black plexiglass (Fig. 10Ai), with odours delivered through an odour port located in the bottom third of one of the short side and connected to the olfactometer output. Infrared top cameras were used to record their behaviour. Following placement in the behavioural chamber, mice were habituated to the arena for 5 minutes, before being exposed to five 60s presentations of solvent separated by 60s intertrial intervals (Fig. 10Aii), followed by three 60s presentations of 1% MT. After habituation to repeated solvent presentations, investigation time withing 3cm of the odour port increased significantly upon first exposure to MT compared to the final solvent presentation (Fig 10Aiii; 5th solvent: 0.85 ± 0.50s; 1st MT: 1.90 ± 0.85s; mean ± SEM; paired t-test, t(7) = −2.68, p = 0.0315). This increase in investigation indicates that mice detected and habituated to the solvent, and then discriminated the odour delivered by the olfactometer from the solvent (36).

**Fig 10.**
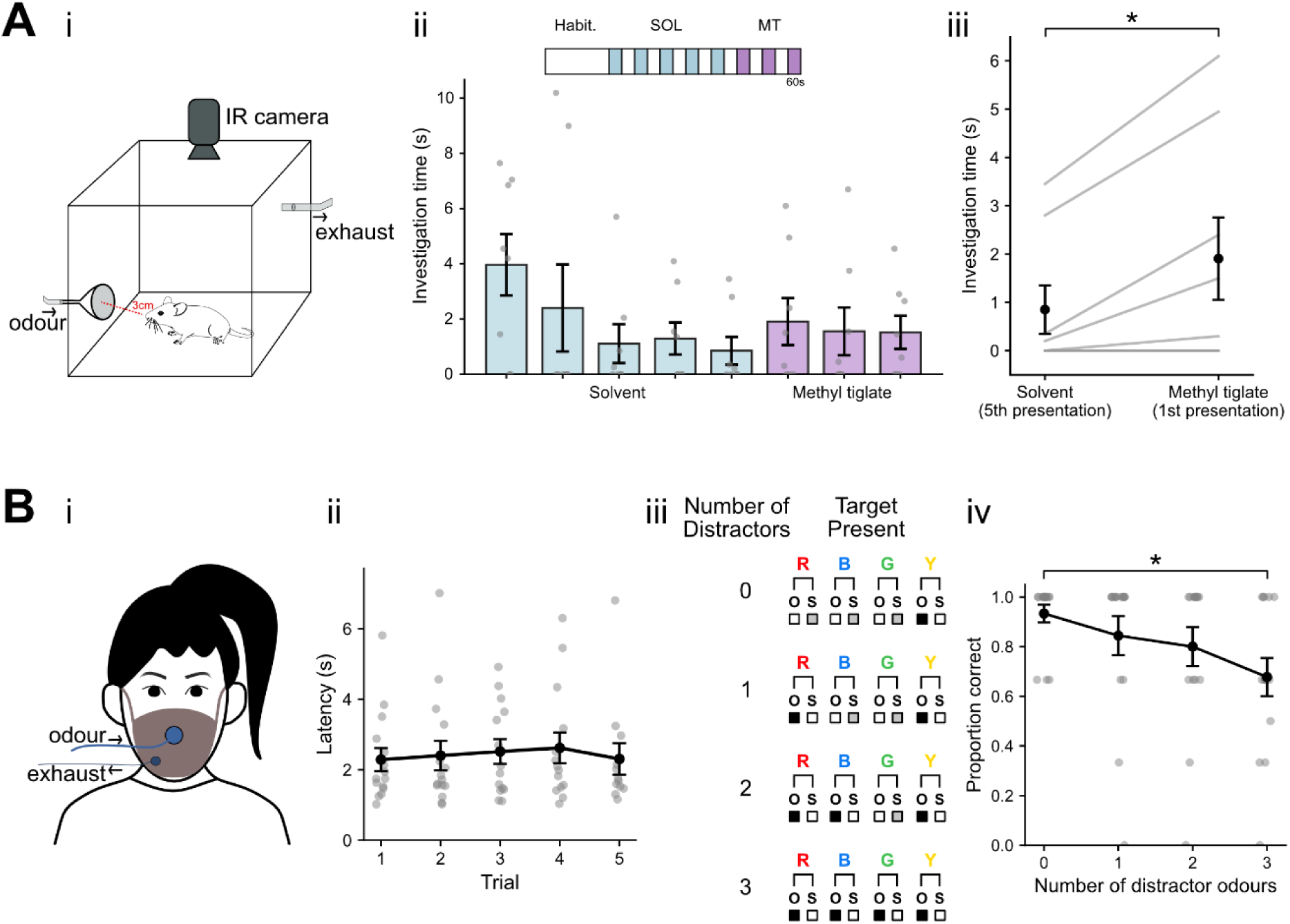
Biological validation of the olfactometer. **(Ai)** Experimental apparatus for rodent testing. Mice were placed in a behavioural arena with an odour delivery port and an exhaust fan to remove odours. IR cameras were used to record behaviour and track investigations (mouse nose within 3cm of odour port). (Aii) Above: Experiment protocol; the coloured blocks represent 60s of solvent or MT delivery, with 60s intertrial interval. Below: Bar plots of odour port investigation duration of one example animal across the whole experiment. (Aiii) Odour investigation times during fifth solvent presentation and first MT presentation. Data in black displays mean ± standard error of the mean; grey lines represent individual animals, n=8. *p *<* 0.05. **(Bi)** Experimental apparatus for human testing. Subjects were fitted with a disposable FFP2 mask fitted with an odour port and an exhaust port. Readout was via a computer keyboard. (ii) Detection latency across repeated odour presentations. (iii) Mixtures experiment protocol. (iv) Detection accuracy across increasing numbers of distractor odours. Data in black displays mean ± standard error of the mean; grey lines represent individual participants, n=15. *p *<*0.05. Grey lines or circles are individual subjects, black circles and lines are mean ± SD.

#### Human behaviour

It is well-known that humans become worse at identifying the presence of particular odours as the total number of odours in an odour mixture increases (37–39). Similarly, performance at identifying a target odour declines as the number of distractor (*i.e.,* non-target) odours increases (40). To validate our custom olfactometer both in terms of reliable delivery of complex odour mixtures and its suitability for human psychophysics, we sought to replicate these established effects. To do this, we conducted two tasks measuring odour detection latency for single odour presentation, and target odour detection with mixtures of increasing complexity.

In the first experiment, participants were presented with five 10s presentations of 2% citronellal and instructed to report odour detection as quickly as possible by pressing the spacebar on a computer keyboard. Participants reliably detected the citronellal delivered by the olfactometer with short and consistent reaction times (Fig. 10Bii). Mean detection latency was 2.43 ± 0.17s (mean ± SEM; n = 15 participants), confirming rapid perceptual detection. Detection latency remained stable across repeated presentations, indicating reproducible stimulus timing and consistent odour delivery across trials.

In the second experiment, participants performed a target odour detection task in which they reported the presence of a target odour (2% citronellal, previously learned during the latency experiment) presented alone or in combination with one, two, or three distractor odours (Fig. 10Biii). Participants reliably detected the target odour, but detection accuracy decreased significantly as the number of distractors increased (Fig. 10Biv). A generalised linear mixed-effects model revealed a significant negative effect of distractor number on detection accuracy (*β* = −0.577, SE = 0.203, *z* = −2.85, *p* = 0.0044), indicating that each additional distractor reduced the likelihood of correctly detecting the target odour.

Thus by replicating previous findings (39,40), these results validate the olfactometer as a reliable tool for precise delivery of single odours and odour mixtures, and for and psychophysical measurement in humans. Notably, the olfactometer has been validated with output tube lengths of up to 4 m (Fig. 5C) and could likely be extended further with appropriate compensation for increased odour delivery latency (supplementary figure 3). This makes it potentially suited for use in combination with neuroimaging experiments, such as fMRI, where metal components require the device to be located outside the scanner room.

## Discussion

Existing olfactometer designs provide reliable odour delivery but often involve trade-offs between cost, flexibility, and ease of use. Here, we developed a modular, low-cost olfactometer to address these limitations while maintaining precise and reliable performance. In designing this olfactometer, we aimed to balance accessibility, flexibility, and experimental precision. The system was developed to be affordable, modular and scalable, and simple to assemble without specialised engineering expertise, while supporting a range of experimental paradigms including rodent behavioural studies and human psychophysics.

Control over odour delivery is implemented using fully open-source software, allowing odour delivery protocols to be easily customised. The olfactometer is built from commercially available components and costs, as of September 2025, approximately £1500/€1700/$2000 (excluding the air source and control computer). Although described here as a four-odour-module system, the modular architecture allows straightforward expansion to additional channels if required. Extensive validation demonstrated highly reliable odour delivery with minimal trial-to-trial variability and consistent performance across modules. The system produced stable signals across stimulus durations ranging from hundreds of milliseconds to several minutes. Together with behavioural validation in both mice and humans, these results indicate that the olfactometer provides precise and reliable odour delivery suitable for diverse experimental applications.

An additional advantage of the design is its ability to deliver odour mixtures while maintaining constant airflow and pressure (21,41). Maintaining stable pressure is critical for mixture experiments, as fluctuations in airflow can confound behavioural and physiological responses. By ensuring constant pressure across conditions, the system enables reliable presentation of complex odour mixtures and supports experiments investigating olfactory processing in more naturalistic sensory environments.

### Limitations

Relative to purchasing a commercial olfactometer, an open-source system requires additional effort to source components, assemble the device, and verify performance in the intended experimental configuration. For groups with limited time but sufficient financial resources, a commercial system may therefore be more practical. Conversely, laboratories prioritising flexibility, modularity, and cost-effectiveness may benefit from the additional upfront investment required for assembly and characterisation.

Although odour delivery dynamics in our current configuration are suitable for many behavioural and physiological paradigms, experiments that require highly complex or rapidly varying concentration waveforms (*e.g.,* sinusoidal modulation or other precisely shaped stimulus profiles) may still benefit from specialised commercial hardware or more sophisticated designs (42). Users implementing the system in new contexts should therefore validate stimulus timing and concentration dynamics under their specific flow rates, tubing lengths, and experimental set ups.

Auditory cues from solenoid valve actuation remain a potential confound in behavioural tasks, particularly learning paradigms. These cues can be mitigated by increasing physical separation between the device and the testing area, acoustically damping the enclosure, and using masking noise (as in our human experiments). Where necessary, probe trials in which odour identities are dissociated from valve patterns can be used to confirm that behaviour is guided by olfactory rather than auditory information (43).

Finally, as with most olfactometers, odour carryover and adsorption to tubing cannot be fully eliminated. Regular replacement of tubing and routine maintenance are therefore recommended, particularly when switching between strongly adsorbing odorants or experimental series. We additionally recommend that users validate output stability after any substantial modification (*e.g.,* tubing length, flow settings), as these factors can influence delivered concentration profiles.

### Improvements and alterative designs

Although the design presented here provides reliable and flexible odour delivery, several modifications could further improve performance depending on experimental requirements.

First, tubing materials could be upgraded. While the current tubing provides adequate performance for most experiments, PTFE exhibits lower odour adsorption and may further reduce contamination between stimuli, albeit at extra expense.

Second, the flowmeters used to regulate airflow could be replaced with mass flow controllers. These devices provide more precise and programmable flow regulation, potentially improving concentration stability and facilitating automated control of airflow parameters. However, mass flow controllers are substantially more expensive and therefore may not be justified for laboratories seeking a low-cost solution.

Third, the electronic control system could be implemented using printed circuit boards (PCBs) rather than the manually assembled boards described here. PCBs would likely increase reliability and simplify wiring once produced, although their construction requires slightly greater familiarity with soldering and electronic assembly. Alternative valve configurations could also be adopted. For example, 3-way solenoid valves could replace the 2-way valves used in our present design (21). This configuration would simplify the electronics required to control the valves (Fig. 2), which may be advantageous for some users. Finally, for users requiring shorter odour onset times, the design could be modified to incorporate a four-way valve near the stimulus outlet. In this configuration, carrier air continuously flows through the odour vial, with the odourised stream diverted to an exhaust under baseline conditions. Upon valve actuation, this stream is redirected to the output, enabling faster odour delivery. However, such configurations may increase the risk of odour depletion during prolonged stimulation and therefore require careful validation.

## Conclusion

We have aimed to provide a detailed description of our olfactometer design, such that it can serve as a practical reference for researchers wishing to build similar systems. The system is affordable, uses readily available components, and enables precise odour delivery. While improvements are certainly possible, we hope this work helps lower technical and financial barriers to olfactory research and encourages broader investigation of this fundamental sensory modality in both humans and animal models.

## Materials and Methods

### Odorants

Odorants were obtained from Sigma-Aldrich (Methyl butyrate, 246093; Pentyl acetate, 46022; Eugenol, E51791; 2-Phenylethanol, 77861), Tokyo Chemical Industry (Methyl tiglate, T0248; Ethyl valerate, V0004), or Thermo Fisher Scientific (Citronellal, 405291000). Odorants were diluted to the desired concentration in diethyl phthalate solvent (Sigma-Aldrich, 524972). All odour concentrations are reported as volume/volume (v/v). Dilutions were prepared immediately before each miniPID recording or behavioural experiment.

### Odour delivery

In all miniPID and behavioural experiments air was supplied via a medical air compressor (MGF compressors, S-OF100-003-MS2). Air was passed through two pressure regulators (one for carrier line, one for odour/regulator lines) set to 0.1MPa. Carrier path flowmeter was set to 1L/min and the regulator and four odour path flowmeters were set to 0.1/min. Odour delivery was controlled using an Arduino microcontroller board (Arduino Mega 2560 R3) programmed with a custom Arduino IDE script. Digital output pins from the microcontroller were interfaced with valve driver boards to precisely control valve activation timing and duration.

### Odour delivery validation and pressure recordings

Odour delivery was confirmed and quantified using a mini photoionization detector (miniPID 200C; Aurora Scientific) positioned at the odour port of a behavioural arena. The miniPID was allowed to warm up for at least 30 minutes before recording. The miniPID was not calibrated using a reference gas with a known absolute concentration, as such, the output voltage reflects only the relative odour concentration rather than an absolute value. Valve openings were simultaneously recorded by monitoring the digital output voltage (0V/5V) from an Arduino microcontroller (Arduino Mega 2560 R3) via a Digidata 1332A (Axon Instruments) and captured with Clampex 9.2 (Molecular Devices). MiniPID recordings were baseline-corrected using a pre-stimulus window.

Odour onset latency was defined as the time from valve actuation to the first increase in the derivative of the signal exceeding 3 × standard deviations (SD) of baseline, following Savitzky–Golay smoothing. Peak amplitude was defined as the maximum baseline-corrected signal during valve opening. Area under the curve (AUC) was calculated over the response period. Decay time was defined as the time from valve closure to when the signal returned to less than 10% of peak amplitude. Pressure was monitored with a digital manometer (Danoplus DP-103), with readings extracted from video recordings of valve operation paired with an LED trigger captured by a USB camera. Peak pressure was determined as the highest manometer reading during valve opening.

### Mouse behavioural testing and analysis

Mice of either sex, aged 2 to 4 months, were housed under a 12 h light/dark cycle in an environmentally controlled room with ad libitum access to food and water. In line with the 3R principles, we used both wild-type mice (C57Bl/6J; Charles River Laboratories) and surplus heterozygous animals from transgenic breedings (R26-Gcamp3 (Gt(ROSA)26Sortm38.1(CAG-GCaMP3)Hze/J; Jax stock #014538) and TRAP (Fostm2.1(icre/ERT2)Luo/J; Jax stock #030323) ongoing in the laboratory. The integrity of the transgenic lines was ensured by generating breeders via back-crossing heterozygous carriers with C57Bl6 animals specifically bought biannually from Charles Rivers Laboratories. All experiments were performed at the University of Cambridge in accordance with the Animals (Scientific procedures) Act 1986 and with AWERB (Animal Welfare and Ethical Review Board) approval.

Mice were placed in behavioural arenas and habituated to carrier airflow for 5 minutes before testing. Odour delivery consisted of five 1-min presentations of solvent, each separated by 60s intertrial intervals, followed by three 1-min presentations of 1% MT [30,39]. Valve timing was controlled by an Arduino microcontroller (Arduino Mega 2560 R3), and odour on periods were signalled by an LED light visible to the camera. Mice were tested in a 30×20×30cm arena made of black IR-transmitting plexiglass. An odour port was installed 5cm from the base of one wall to deliver odour. An exhaust tube on the opposite wall removed residual odour via an extractor pump (Charles Austen Pumps Ltd, LD10).

Behaviour was recorded at 20fps using an IR USB camera top-down using Bonsai-Rx (44). All testing was conducted in darkness or red light. Body tracking was performed in DeepLabCut(45), using the superanimals topviewmouse model from Modelzoo (46). The data from seven body parts were extracted and fed into SimBA (47) for behavioural quantification. A circular region of interest (ROI) with radius 3cm was drawn centred on the odour port. Investigation time was calculated by taking the time spent with the nose located within the ROI. Statistical analyses were performed using RStudio (version 4.5.0). Data normality was assessed using the Shapiro-Wilk test, with parametric or nonparametric tests applied accordingly. Statistical significance was set at *α* = 0.05 for all two-tailed comparisons.

### Human behavioural testing and analysis

Fifteen healthy adult volunteers of either sex participated in the human olfactory experiments. Participants were not pre-screened for olfactory function. Written informed consent was obtained from all participants prior testing, and no personally identifiable information was collected. All procedures were conducted at the University of Cambridge and were approved by the Cambridge Human Biology Research Ethics Committee (approval number HBREC.2025.29).

Odours were delivered using the custom-built olfactometer controlled by an Arduino microcontroller (Arduino Mega 2560 R3) and custom-written Python scripts. Participants wore an FFP2 mask fitted with two ports: one connected to the olfactometer output tubing (70 cm) and the other to an extractor pump (Charles Austen Pumps Ltd, LD10) to prevent odour accumulation. To minimise auditory cues from valve actuation, participants wore noise-cancelling headphones playing music. Prior to testing, participants were familiarised with all odours at the concentrations used in the experiment.

In the latency experiment, participants were seated in front of a computer and received five 10s presentations of 2% citronellal separated by 15s intertrial intervals. Participants were asked to press the spacebar upon detecting the odour. Detection latency was defined as the time from odour onset to response. Trials without a response within the 10s response window were excluded from the latency analysis (4/75 trials). In the demixing experiment, participants performed an odour detection task in which they were required to report the presence of a target odour (2% citronellal) presented alone or in mixtures with one, two, or three distractors (eugenol, 2-phenylethanol, and ethyl valerate; all 2% concentration). On each trial, four olfactometer channels were activated, with non-odour channels delivering solvent to maintain constant airflow. Target-present trials included four conditions (target only, target + 1, 2, or 3 distractors), and target-absent trials comprised the corresponding distractor-only conditions, yielding eight conditions in total (four target-present, four target-absent). Each condition was repeated three times. Distractor identity in the one- and two-distractor conditions was counterbalanced across participants by assigning them to one of three groups, while all participants received all distractors in the three-distractor condition. Each stimulus was presented for 15s with 15s intertrial intervals. Participants responded using a keyboard to indicate whether or not the target odour was present. Detection accuracy was defined as the proportion of correct responses on target-present trials. Accuracy was analysed using a generalized linear mixed-effects model with a binomial error distribution and logit link function implemented using the lme4 package in RStudio (version 4.5.0). Distractor number was included as a fixed effect and participant as a random intercept to account for repeated measures. Statistical significance was set at *α* = 0.05.

## Acknowledgements

This work was supported by project grants from the Newton Trust, the URKI Biotechnology and Biological Sciences Research Council (BB\W014688\1), and the Wellcome Trust (301427/Z/23/Z) (EG); a Homerton College Junior Research Fellowship (CG); and a Harding Distinguished Postgraduate Scholarship (CD). We wish to thank all of the participants who volunteered to help in our study; Dan Rokni, Tobias Ackels, Jasper Poort, Tom Oldham, and Glenn Harrison for their helpful insights during the designing and validating stages; Harin Wijayathunga and Sonu Peedikayil Kurien for beta testing the construction instructions; Edina Horvath-Gulasci for help running the mouse experiments; Ailie McWhinnie for comments on the manuscript; participants of European Chemoreception Research Organization 2025 conference and all members of the Galliano laboratory for helpful discussions.

## Supporting information

**S1.**
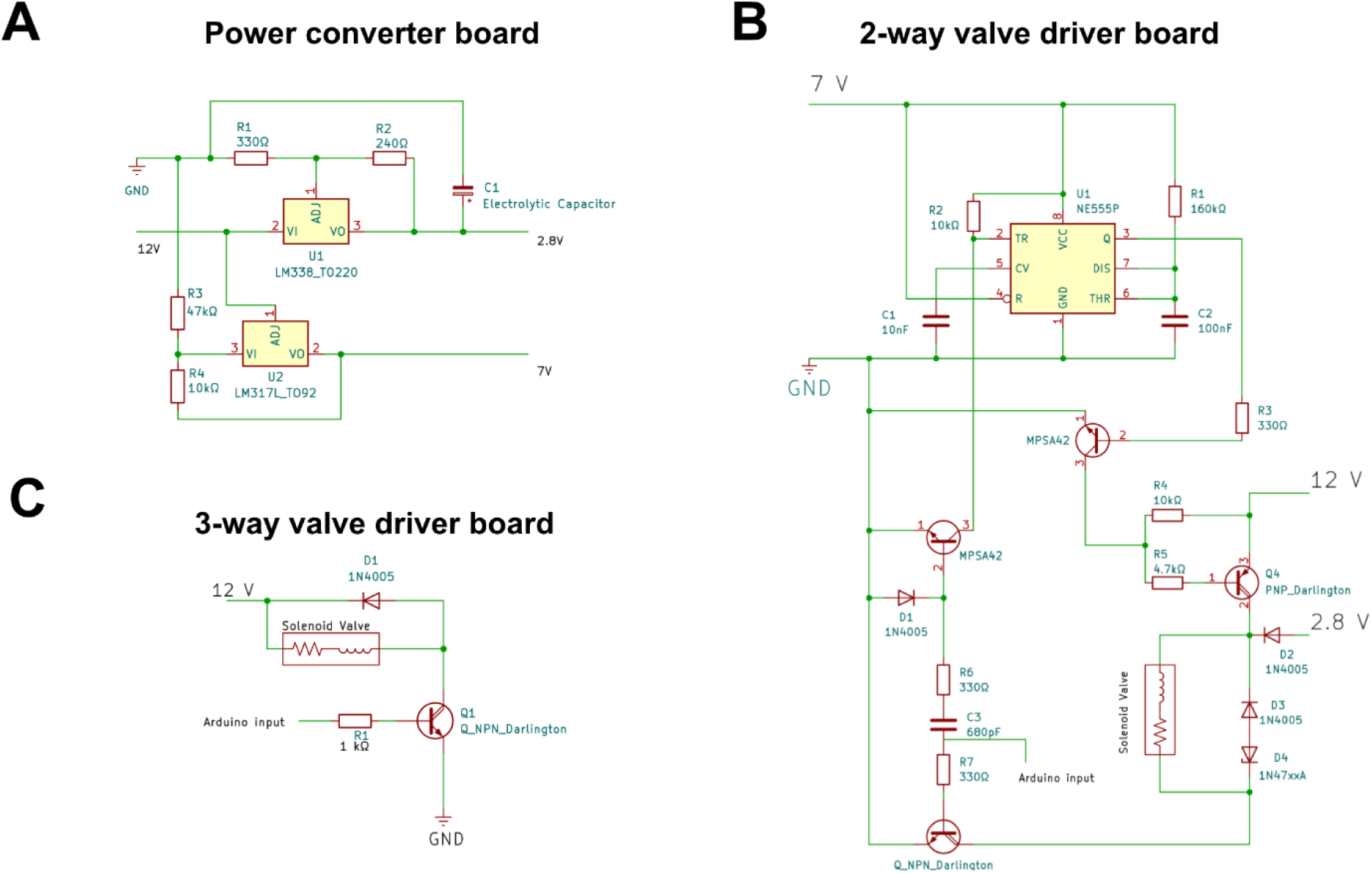
Electronic schematics of the circuits used to control the olfactometer. **(A)** Power converter board circuit. **(B)** 2-way valve driver board circuit (shown for a single valve; two of these circuits are implemented on the breadboard in Fig 2). **(C)** 3-way valve driver board circuit. Related to Fig. 2.

**S2.**
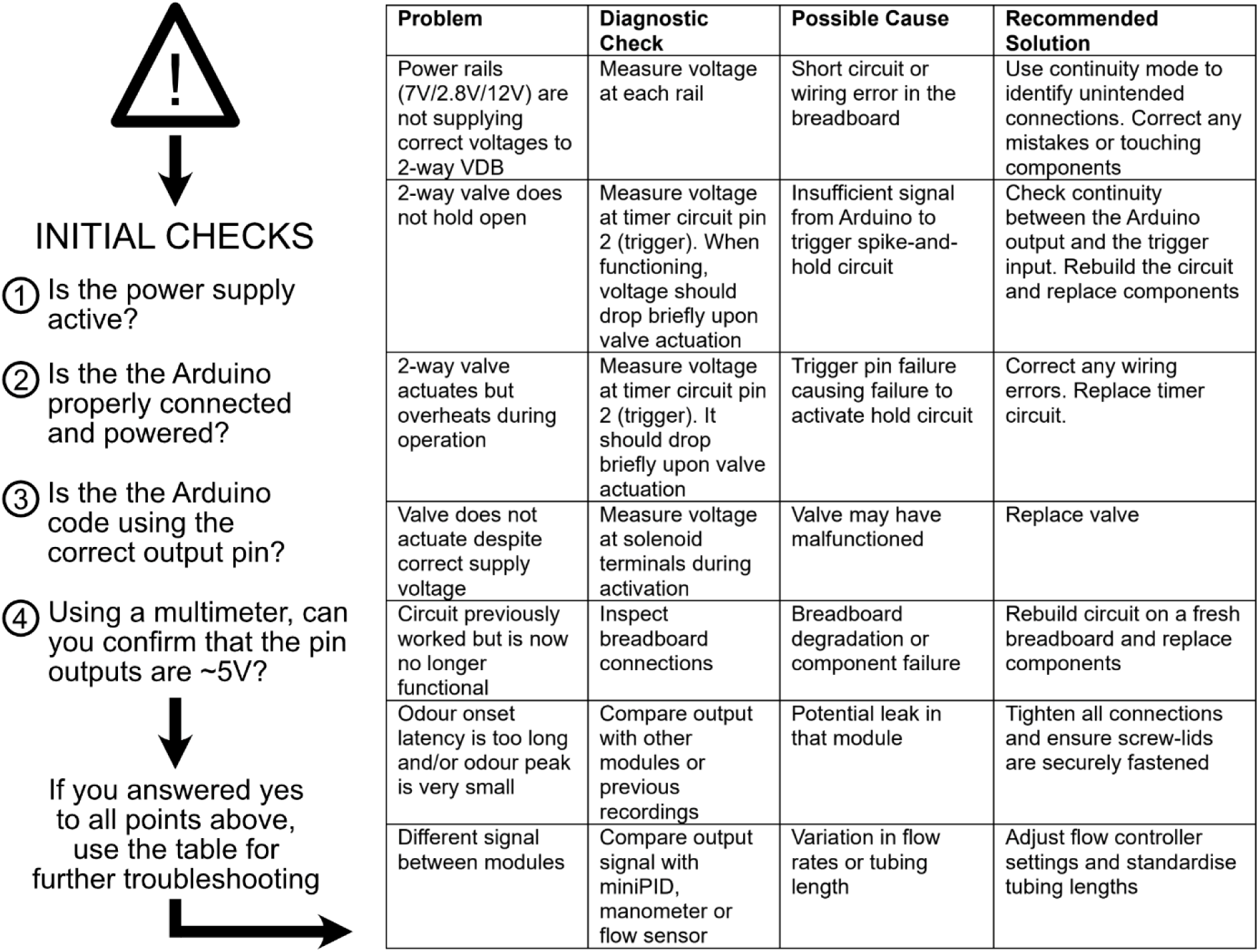
Troubleshooting workflow and tips.

**S3.**
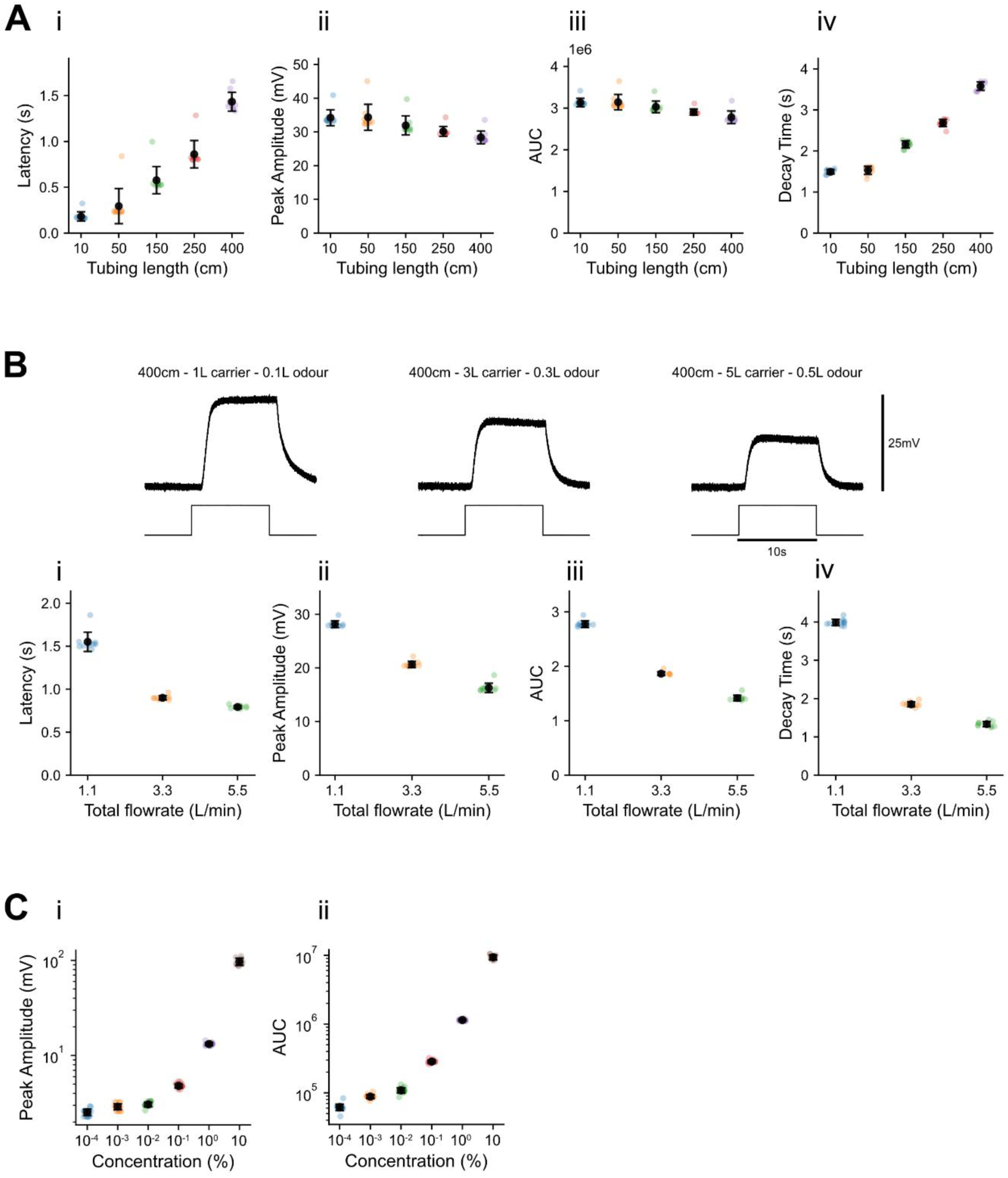
Quantitative assessment of the influence of tubing length, total flow rate, and liquid dilution on recorded signals. **(A)** Quantified features of 5% MT delivered across tubing lengths ranging from 10–400 cm (Fig. 5), including (i) latency to odour onset, (ii) peak amplitude, (iii) area under the curve during valve opening, and (iv) decay time. **(B)** Example miniPID trace during a 10s exposure of 5% MT under varying carrier and odour flow rates with 400cm tubing length (Fig. 5), in which the total flow rate was increased while maintaining a consistent odour-to-carrier ratio. Top: miniPID signal; bottom: digital output voltage trace from the Arduino indicating valve opening. Carrier and odour flow rates are indicated above each trace. Quantified features include (i) latency to odour onset, (ii) peak amplitude, (iii) area under the curve during valve opening, and (iv) decay time. **(C)** Quantified features of MT delivered across a concentration range from 10-4% to 10% (Fig. 7C), including (i) peak amplitude and (ii) area under the curve during valve opening.

**S4.**
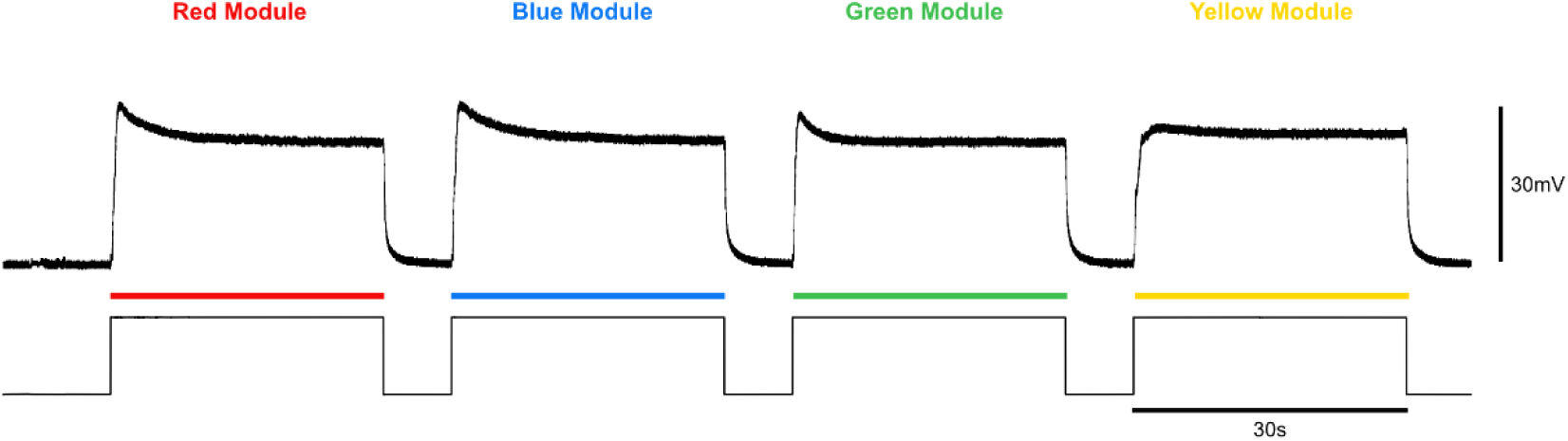
Extended odour delivery across modules. Recording trace of sequential 30s presentations of 5% MT from all four modules (related to Fig. 8). Top: miniPID signal; bottom: digital output voltage trace indicating valve state.

**S5.**
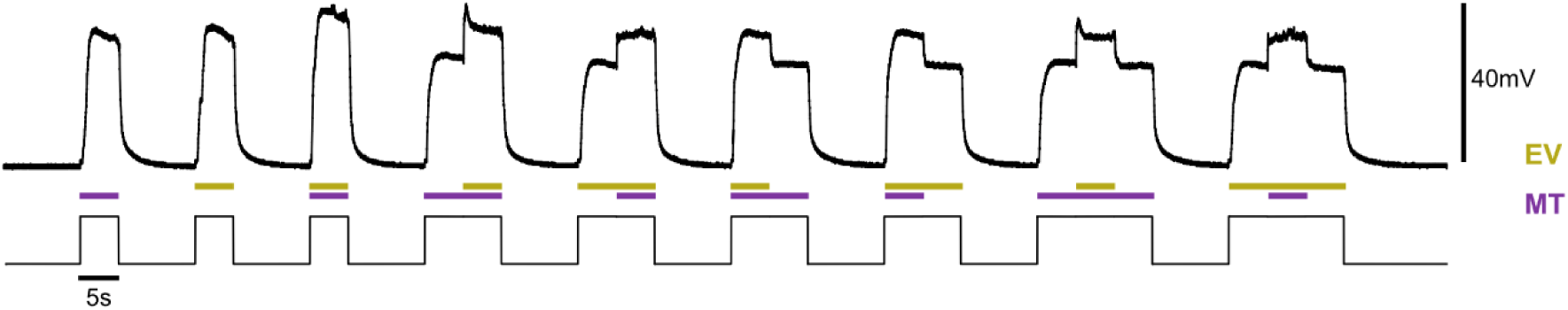
Flexible timing of odour mixtures. Recording trace of 5% EV and 5% MT presented alone and in combination, showing sequences of single-odour and mixture periods with different timing of delivery (related to Fig. 9) Top: miniPID signal; bottom: digital output voltage trace indicating valve state.

## Notes

### Competing Interest Statement

The authors have declared no competing interest.

### Summary of Updates

The overall design presented in the manuscript has been updated. The new manuscript reflects these changes.

